# Encoding of 2D self-centered plans and world-centered positions in the rat frontal orienting field

**DOI:** 10.1101/2022.11.10.515968

**Authors:** Liujunli Li, Timo Flesch, Ce Ma, Jingjie Li, Yizhou Chen, Hung-Tu Chen, Jeffrey C. Erlich

## Abstract

The neural mechanisms of motor planning have been extensively studied in rodents. Preparatory activity in the frontal cortex predicts upcoming choice, but limitations of typical tasks have made it challenging to determine whether the spatial information is in a self-centered direction reference frame or a world-centered position reference frame. Here, we trained rats to make delayed visually-guided orienting movements to six different directions, with four different target positions for each direction, which allowed us to disentangle direction versus position tuning in neural activity. We recorded single unit activity from the rat frontal orienting field (FOF) in the secondary motor cortex, a region involved in planning orienting movements. Population analyses revealed that the FOF encodes two separate 2D maps of space. First, a 2D map of the planned and ongoing movement in a self-centered direction reference frame. Second, a 2D map of the animal’s current position on the port wall in a world-centered reference frame. Thus, preparatory activity in the FOF represents self-centered upcoming movement directions, but FOF neurons multiplex both self- and world-reference frame variables at the level of single neurons. Neural network model comparison supports the view that despite the presence of world-centered representations, the FOF receives the target information as self-centered input and generates self-centered planning signals.

## Introduction

We use multiple reference frames to represent the world. For example, as you plan a movement to reach for your morning coffee, the arm region of motor cortex may represent the goal in an arm-centered reference frame and your frontal eye field represents the goal in an eye-centered reference frame. Other areas of your brain may represent the cup relative to the room or table or the milk or sugar. During sensorimotor behaviors, sensory information is initially represented in sensor reference frames and motor commands are finally represented in muscle reference frames. Since the sensors and effectors are embodied, we can think of these representations as being in self-centered (or egocentric) reference frames, i.e. they are reference frames that move around with the subject as they move through the world. However, our lived experience is in a world-centered (or allocentric) reference frame: we feel as if we move around and make decisions in a stable world. Moreover, allocentric representations are found in a wide range of brain regions (Hafting et al., 2005, O’Keefe and Nadel, 1978, Taube et al., 1990, Wilber et al., 2014). Thus, a full understanding of the neurobiology of motor planning needs to address the question of where and how these reference frame transformations take place (Andersen et al., 1990, 1985, Andersen and Mountcastle, 1983, Cohen and Andersen, 2002).

The neural mechanisms of motor planning in rodents have been extensively studied in two-alternative forced choice (2AFC) and go-nogo tasks (Chen et al., 2017, Erlich et al., 2011, Guo et al., 2014, Sul et al., 2011). Converging evidence has implicated the frontal orienting field (FOF), a subregion of the secondary motor cortex (M2), as a cortical substrate for planning orienting movements (Erlich et al., 2011, Hanks et al., 2015, Olson et al., 2019), especially when those plans require flexible sensorimotor processes (Erlich et al., 2015, Siniscalchi et al., 2016, Zhu et al., 2021). Quantitative models suggest that the FOF is a part of a bistable attractor network for short-term memory and decision making in 2AFC orienting movement planning (Hanks et al., 2015, Kopec et al., 2015, Piet et al., 2017). Similar work in the mouse anterior lateral motor cortex (ALM) during directional licking also identified discrete bistable attractor models as best accounting for observed neural activity and perturbation results (Inagaki et al., 2019, Li et al., 2016). Bistable attractor models are currently state-of-the-art for 2AFC motor planning, but they are ambiguous as to the spatial reference frame of the neural coding. Do they represent the planned movement direction in an self-centered reference frame, the target location in an world-centered reference frame, or a ‘decision’ in an abstract reference frame? Given the limitations of those behavioral paradigms, the answer is largely unknown.

The FOF is a potential site to integrate egocentric and allocentric spatial representations, as it receives input from the posterior parietal cortex (Reep et al., 1994) and the retrosplenial cortex (Yamawaki et al., 2016), both of which exhibit egocentric as well as allocentric spatial representations (Wang et al., 2020). To test the reference frame of action planning representation in the FOF, we designed a multi-directional, multi-positional orienting task that could distinguish between allocentric and egocentric reference frames. We found preparatory and movement-related activity in the FOF that were in the egocentric reference frame. Interestingly, allocentric target position information was also encoded in the FOF, but emerged later than direction encoding. The allocentric encoding represented the *current* rather than the upcoming position of the animals, indicating the allocentric activity did not play a primary role in action planning. These two reference frames were multiplexed at the single-neuron level: task-related activity was best described as an egocentric direction tuning multiplicatively modulated by the allocentric current position.

Similar gain fields have been previously reported in primate frontal and parietal eye fields during saccade planning (Andersen et al., 1985, Cassanello and Ferrera, 2010, 2007) and were suggested to support reference frame transformation (Pouget and Sejnowski, 1997, Salinas and Abbott, 1995, Zipser and Andersen, 1988). However, a recurrent network model whose input and output were both in the egocentric reference frame had the most similar activity to the FOF neurons: including representing allocentric activity and gain fields. This finding calls into question the proposed role of gain fields in supporting reference frame transformations. Moreover, it hints that the allocentric representations in FOF are not necessary for the task. These results further support our conclusion that planning in the FOF takes place in an egocentric coordinate frame, although the multiplexed spatial encoding may support spatial information integration in downstream brain areas or online error correction which might be required for planning complex movements sequences.

## Results

We trained rats to perform visually-guided orienting movements to multiple directions and multiple target positions. The training apparatus consisted of a vertical port wall with 7 operant ports and an additional port for reward delivery (Figure 1A). For the majority of sessions, there were 24 trial types: 6 possible directions, with each direction starting from 4 different start positions (Figure S1A). This design allowed us to dissociate self-centered movement direction from the world-centered start and target position (Figure 1B). Throughout the paper we indicate self-centered movement directions with a blue-red (left-right) and dark-light (down-up) color scheme (Figure 1C). For world-centered positions we use a green-orange (left-right) and dark-light (down-up) color scheme (Figure 1D).

**Figure 1.**
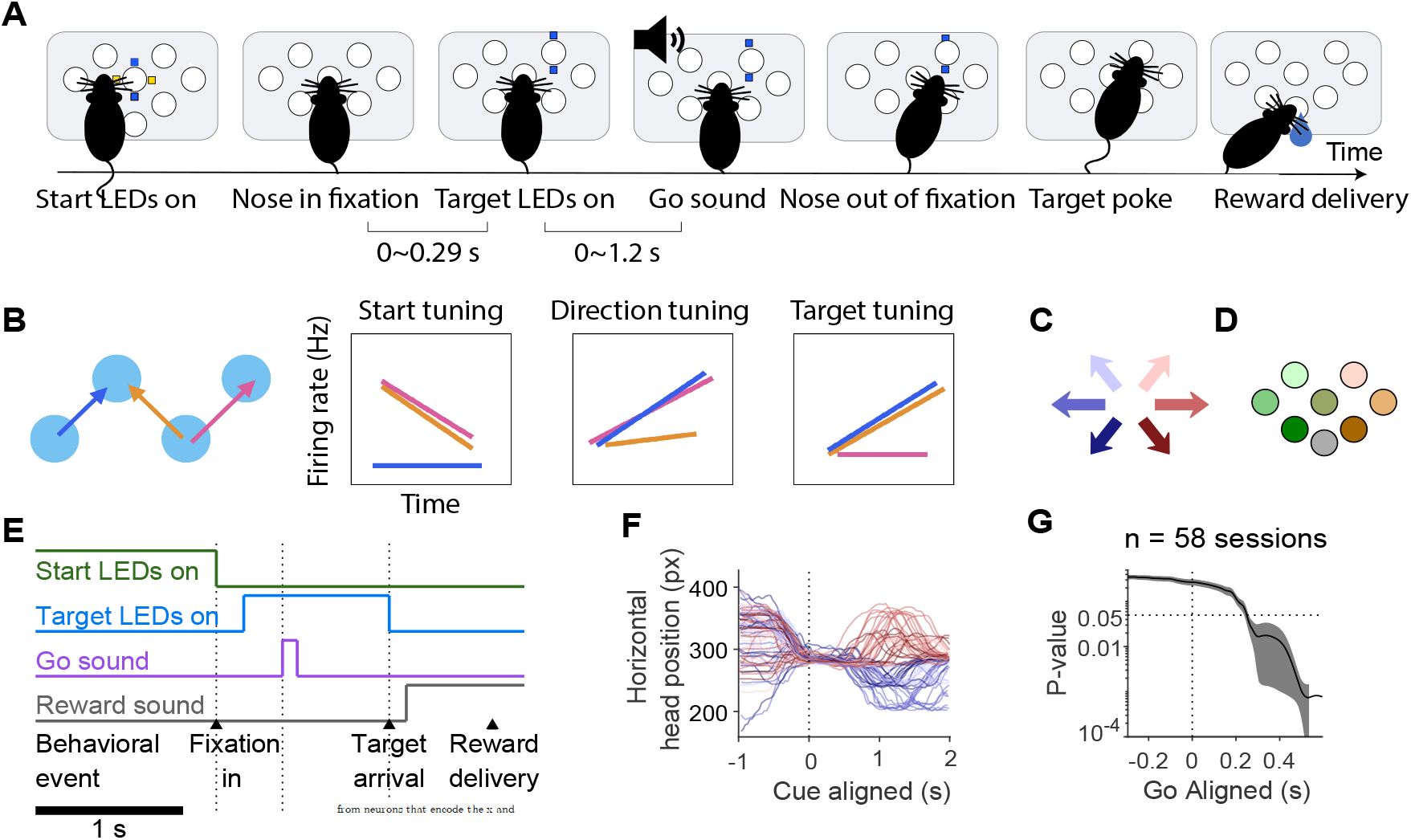
A visually-guided multi-directional orienting task in rats. **A**. Schematic of the task. Each trial began with the onset of a pair of blue and yellow LED, cuing the rat to nose poke into the start port. The start LED extinguished upon arrival in the start port. After a short delay, a blue LED illuminated, indicating the target port. After a go sound, the rat withdrew from the start port and poked into the target port. Water reward was delivered for correctly performed trials. **B**. The logic of the task. We consider three trials and how cells with different task tuning would respond. The orange and pink trials share start position. The blue and pink trials share direction. The blue and orange trials share target. Right, the postulated firing patterns of neurons that were tuned to start position, movement direction, or target position respectively. **C**. The color scheme of the 6 movement directions. **D**. The color scheme of the 7 port positions. **E**. Timeline of a trial in a typical session. **F**. An example of head position tracking data extracted from video using DeepLabCut. Each line is the horizontal head coordinate during a trial, in unit of pixels from the video frames. Only trials starting from the central port are shown. **G**. There is no correlation between the horizontal head position and the planned movement direction before the go cue. The line indicates significance level of the movement direction modulating the horizontal head position in unit of pixels across time aligned to the go sound, where the effect of start position was captured in the random effect. Line and error bars, mean *±* s.d. of the *p* values over the 58 sessions.

Each trial began with illuminating the yellow and blue LEDs around a ‘start port’, which was randomly chosen from one of the 7 operant ports. Rats fixated in the start port until a go sound. The start port LEDs extinguished at the beginning of fixation, and the target port was illuminated with blue LEDs shortly after fixation onset with 0 to 0.29 s delay. For trials that started in the center, one of the six remaining operant ports was chosen at random as the target. For the other start positions, one of three adjacent ports was chosen as the target (Figure S1A). After the go sound, rats withdrew from the start port and moved to the target port. The target port LEDs extinguished once the animal arrived at the target port, or when the animal poked in another port in error. If the rat poked in the correct port, the water delivery port LED illuminated, a “correct” sound was played, and reward could be collected at the reward port (Figure 1A&E). If the rat poked into the wrong port, an “error” sound was played and there was no reward. Animals kept still during the fixation period (Figure 1F&G). A trial was considered incomplete if the animal did not poke into any port after the start port within 15 seconds. Unless otherwise specified, incomplete trials and fixation violations were not included in analyses. Rats performed 318.89 *±* 7.76 (mean *±* s.e.) trials in each 1.5 hour recording session. As expected from a visually-guided task, performance was good (% Correct = 94.05% *±* 0.53%, mean *±* s.e., n=104 sessions; Figure S2A).

### FOF neurons were tuned to self-centered movement directions and world-centered head positions

We recorded 1224 single neurons in the FOF from 104 sessions in 4 rats. Consistent with previous findings, there were neurons tuned to the upcoming and ongoing movement (Figure 2). To quantify the relative strength of tuning to the start position, direction or target position in single neurons, we fit spike counts in 500 ms time windows to three Poisson generalized linear models (GLMs), whose independent variables were the start position, movement direction or target position, respectively. The spatial variables were coded as factors to avoid assuming any specific functional form of tuning. As we expected, there were neurons more strongly tuned to the self-centered movement direction than the world-centered target position, during the planning phase (Figure 2A-C) or the execution phase (Figure 2D-F,G-I). Interestingly, there were also neurons more strongly tuned to the world-centered target position than self-centered movement directions upon target arrival (Figure 2J-L,M-O,P-R). When firing rates were conditioned on both direction and target position, many neurons seemed to code the conjunction of position and direction (Figure 2C,F,I,O,R). We will address conjunctive coding in later sections (Figure 6); here we will examine another question: which spatial variable do FOF neurons most strongly encode at each phase of the trial?

**Figure 2.**
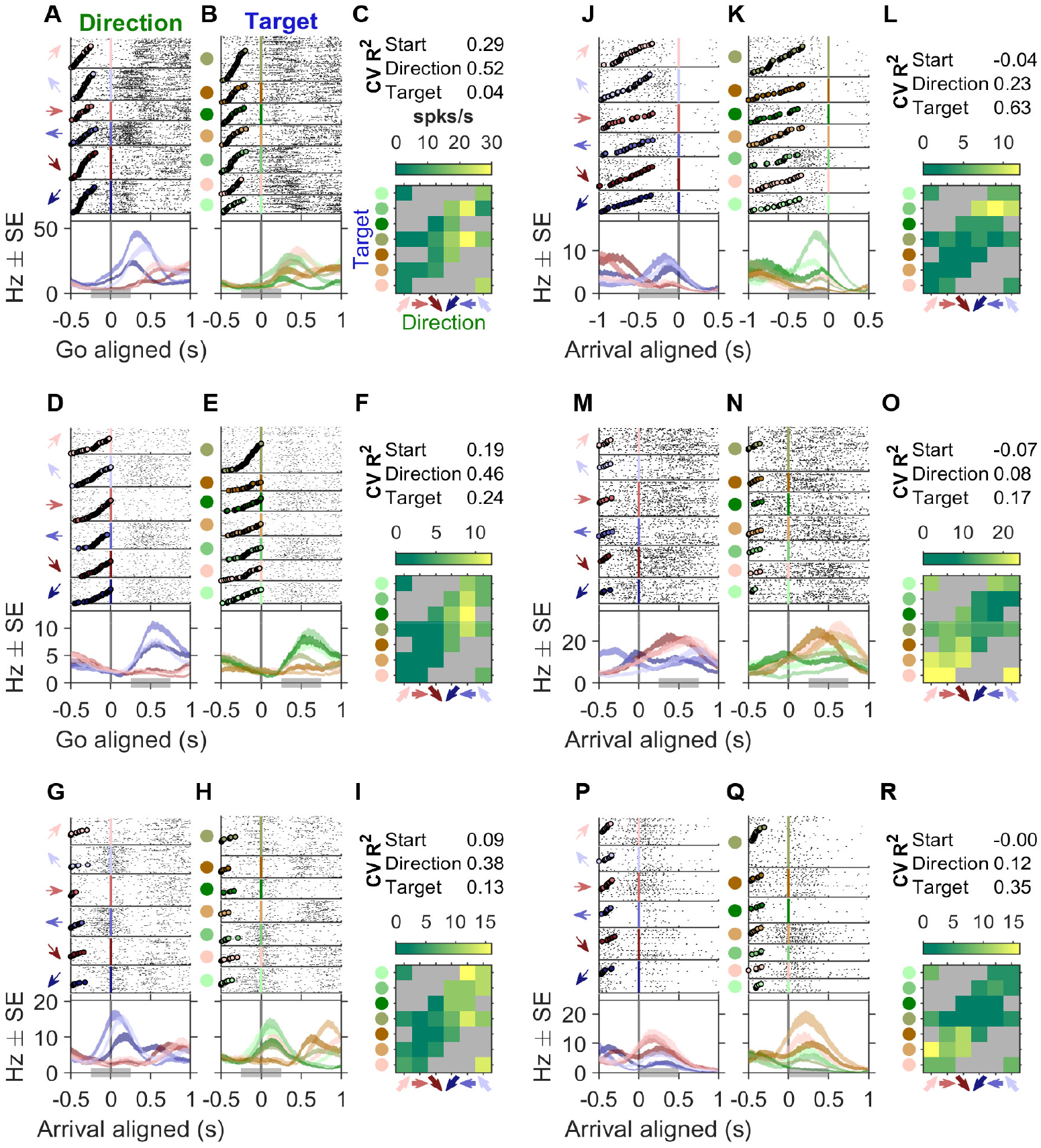
Example neurons with egocentric and allocentric spatial representation. **A-C** An example neuron more modulated by the egocentric movement directions than by the allocentric target positions. **A**. Raster plots and PETHs aligned to the go sound and sorted by movement directions. The top 6 panels show spike rasters grouped by the 6 movement direction, indicated by arrows on the left side of the panels. Circles in each raster panel indicate the time of the visual cue onset on each trial. The bottom panel shows the PETHs of the spikes generated by a causal half-Gaussian kernel with an SD of 200 ms. The shaded areas of the PETHs indicate *±* s.e.. The grey bar at the bottom of the panel indicates the time window to compute the firing rate in each trial type in C. **B**. Raster plots and PETHs of the same cell and same alignment as in **A**, but sorted by target position. Target positions are color coded as in Figure 1D. Circles in each raster panel indicate the time of visual cue onset on each trial. **C**. The maximum *a posteriori* estimated firing rate for each trial type, where the prior was a Poisson distribution whose mean was estimated from all trials. **D-F** and **G-I**, two more example cells as in A-C. **J-L, M-N** and **P-R**, example neurons more modulated by the target position than direction. Neural activity was aligned to the target poke. Circles in each raster panel indicated the time of the go sound.

### FOF neurons encoded self-centered movement plans prior to world-centered target positions

From visual examination of neural activity, there was considerable heterogeneity in task variable tuning as well as the temporal dynamics of tuning across the trial. To get a holistic view of the temporal dynamics of spatial tuning in single neurons, we fit the spike counts of each neuron in four 300 ms time windows to the three Poisson GLMs. The time windows were: ‘pre-cue’, -300 ms to 0 ms aligned to the visual cue onset; ‘post-cue’, 0 ms to 300 ms aligned to the visual cue onset; ‘go’, 0 ms to 300 ms aligned to the go sound; ‘arrival’, -150 ms to 150 ms aligned to the target arrival. Of the 1224 neurons, 541 (44%) were selective to one or more task variables (start position, direction or target position) in at least one time window. We estimated which of the three task variables *best* explained the neural activity in each time window. Most neurons were best tuned to the start position in the pre-cue time window. After the visual cue onset, direction tuning increased, whereas target position tuning emerged even later than movement direction and peaked at the arrival time window (Figure 3A). Thus, the FOF encodes the egocentric movement direction before the allocentric target position, even though the appearance of a visual target cue provided information about the movement direction *at the same time as* the target position. We then extended the analysis by sliding time windows aligned to the visual cue, go sound or target arrival, and the same temporal trend was captured by the *R*^2^s of the corresponding GLMs across time (Figure S7A).

**Figure 3.**
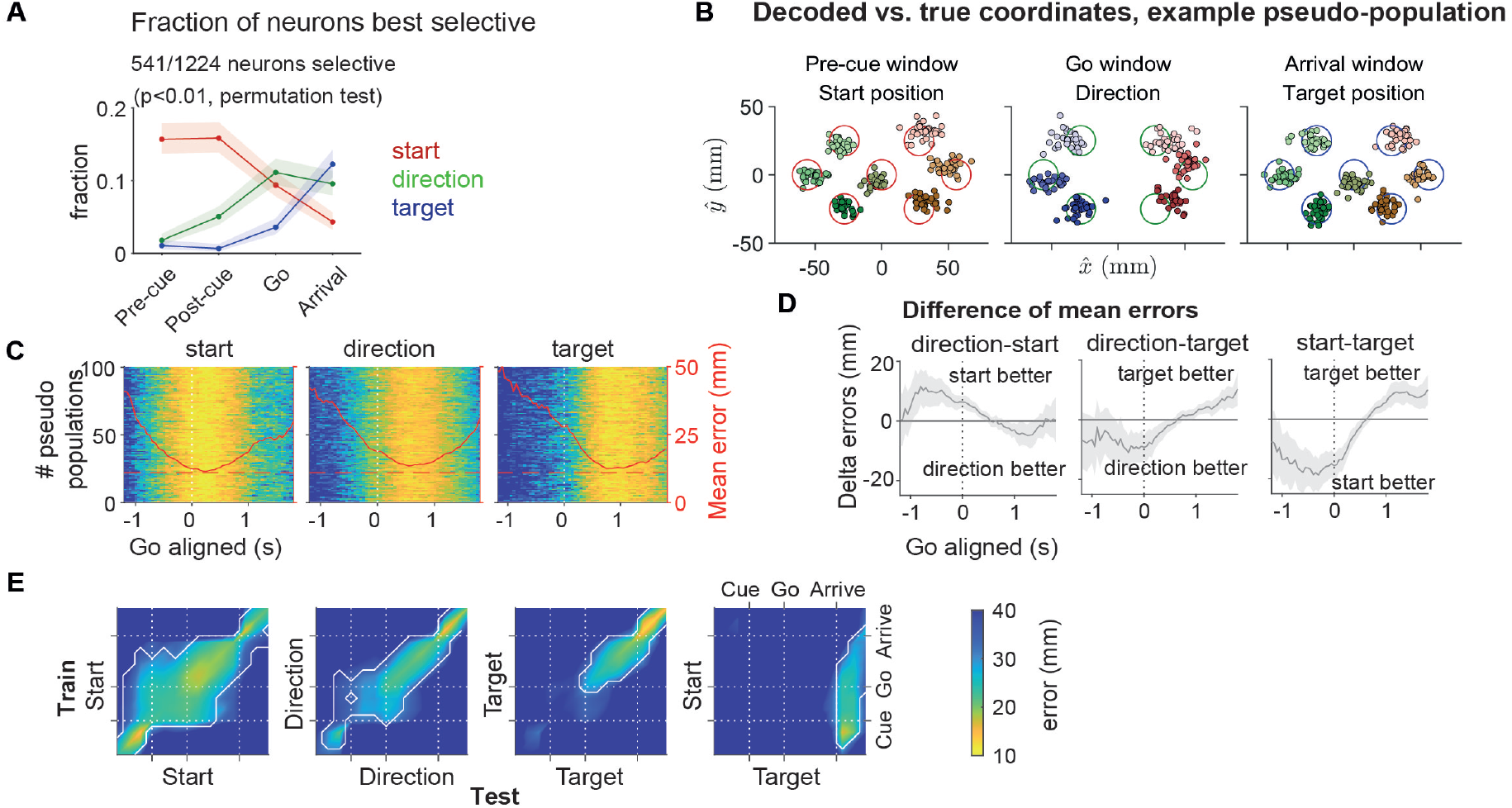
Temporal profile of start position, direction and target position encoding in the FOF population. **A**. The fraction of neurons best selective to the start position (red), the direction (green) or the target position (blue) in 4 time windows (see main text or Methods for definition). Shaded areas indicate the 95% binomial confidence intervals. “Pre-cue”, -300 to 0 ms to visual cue onset. “Post-cue”, 0 to 300 ms to visual cue onset. “Go”, 0 to 300 ms to go sound. “Arrival”, -150 to 150 ms to target poke. **B**. Predicted coordinates of start position, movement vector and target position in an example pseudopopulation decoding using all neurons with enough trials for each type (n = 1194). Each small circle indicates the predicted coordinates in a pseudo trial, where the color indicates the pseudo trial class. Each large circle indicates the coordinates and the diameter of a port on the port wall. **C**. Decoding errors for each pseudopopulation across time aligned to the go sound. Decoding error is defined as the Euclidean distance between the predicted and the actual coordinates, and is indicated by colors following the colorbar in E. Each row is a different pseudopopulation where 1197 neurons were resampled randomly with replacement. Red lines indicate the mean errors across the 100 pseudopopulations. **D**. Mean *±* s.d. of the difference of decoding errors between two spatial variables across the 100 peudopopulations. Positive difference indicates the better decoding of the second variable, and vice versa. **E**. Decoding errors with cross-window decoding. Colors of the heat-maps indicate the mean Euclidean distance between the decoded and true spatial coordinates, averaged across 100 pseudopopulations. The decoders were trained at one time window and tested at another. In the last panel the multivariate linear model was trained for start position and decoded for target position. Contours, *p <* 0.01 (extreme pixel-based test).

In our task, the three task variables were correlated. For example, if the animal started from a port on the left, the movement direction and target position would be likely on the right. As such, a neuron tuned to movement direction could be spuriously found to be tuned for start or target position due to this correlation. To address this potential confound and validate the effectiveness of our method, we generated surrogate spike counts (matched to the tuning and firing rates of real neurons) and found that the errors in classification (e.g. incorrectly labeling a ‘start’ neuron as a ‘direction’ or ‘target’ neuron) to be less than 10% (Figure S7C).

We then used pseudopopulation decoding to examine the geometry of spatial representation on a continuous scale. We pooled all the neurons across sessions where there were at least 8 trials for each start position, direction or target position to construct the pseudopopulation (1197 neurons, 99 sessions, 3 rats). From visual examination, the first 4 principle components of the pseudopopulation activity represented the horizontal and vertical spatial coordinates of the task variables (Figure S8A-F). To prevent a few neurons in the population from dominating the decoding, we randomly resampled the neurons with replacement to construct 100 pseudopulations. We decoded the x and y coordinates of each task variable using multivariate linear decoders with two-fold cross validation, from the first 4 principle components of the pseudopopulation activity in 300 ms time windows (Figure S8G). The errors, defined as the Euclidean distance between the predicted and the actual spatial coordinates, were as small as the radius of the port (around 11 mm) at the best time window for each spatial variable (Figure 3B&C). Put in other words, the geometry of both the movement directions and the port positions were embedded in a linear subspace of the FOF activity.

Pseudopopulation decoding accuracy also demonstrated the sequential encoding of start position, movement direction and target position. The relative goodness of decoding between two spatial variables was quantified as the difference between the two mean decoding errors (Figure 3D). Start position was decoded better than movement direction or target position in the early phase of a trial. At the time of the go cue, direction decoding was significantly better than target decoding. When decoding pseudopopulation activity at one time window with decoders trained at a different time window (Figure 3E), start position tuning was stable across much of the trial. Decoders trained with start positions during fixation could accurately decode target positions around target poke, suggesting a consistent coding for the “current” head position throughout the trial (Figure 3E).

### Single FOF neurons tracked the allocentric current head position

We reasoned that the current position coding in the pseudopopulation could be due to single neurons that had consistent tuning for the current head position (Figure 4A-B). We quantified this consistency using the Pearson correlation between start position tuning in the “pre-cue” window and target position tuning in the “arrival” window, and denoted this as the “start-target tuning correlation” (Figure 4C). Among neurons selective to both the start and the target position (*p <* 0.05 for both the start and the target GLMs, n = 174), the mean start-target tuning correlation was significantly positive (0.66, [0.61,0.70], mean, [95% CI], *p* = 2 *×* 10^*−*4^, permutation test with 10000 replicates) (Figure 4D).

**Figure 4.**
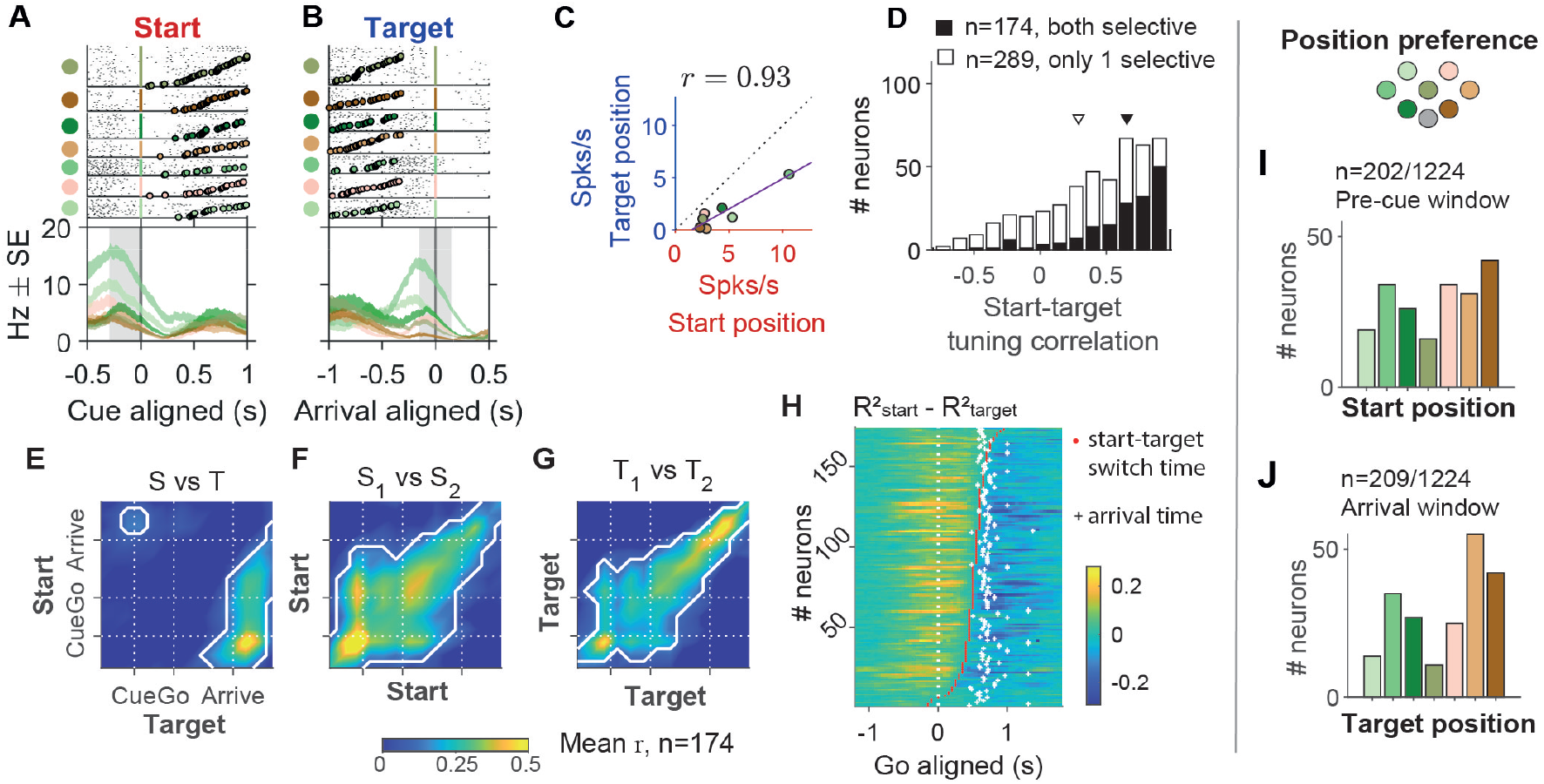
FOF neurons encoded the current head position. **A**. Raster plots and PETHs of an example neuron aligned to the cue, grouped by start position. The shaded grey area indicates the time window to calculate the x-axis firing rate in C. **B**. The same neuron aligned to target poke, grouped by target position. The shaded grey area indicates the time window to calculate the y-axis firing rate in C. **C**. The correlation between start position tuning and y position tuning in the example neuron, where *r* denotes start-target tuning correlation of this neuron as in D. Purple line denotes the total least square fit. **D**. The distribution of the start-target tuning correlation. Black bars are for neurons selective to both start position and target position, and white bars are for neurons selective to only one of the two variables (Mean, [95% CI], 0.66, [0.61, 0.70] for both selective and 0.29, [0.25, 0.34] for only 1 selective). Triangles indicate the means. **E**. The start-target tuning correlation in warped time windows aligned to the visual cue, the go sound and the arrival, averaged across neurons with both start and target selectivity (*n* = 174). The white contours indicate the areas where correlation is significantly larger than 0 (*p <* 0.05 with Bonferonni correction). Different from C and D, these correlations were calculated between start tuning in half of the trials and target tuning in the other half of trials, and vice versa, then took the average. **F**. Similar to E, but for the mean Pearson correlation between pairs of time windows for start position tuning in one half of trials versus the other half. **G**. Similar to F, but for target position tuning. **H**. Time of transition from start position coding to target position coding in single neurons. The color of the heat-map indicate the difference between *R*^2^s of the start position GLM and the target position GLM, calculated at each 300 ms time window aligned to the go sound. Each row was a neuron (n = 174). The red dots indicate the time of switching from *R*^2^ higher to *R*^2^ higher (see Methods for details). The white crosses indicate the averaged time of target poke for that session. **I**. The number of neurons preferring each start position in the “pre-cue” time window, among cells that had significant start position selectivity in the “pre-cue” window (*p <* 0.01, permutation test for the start position GLM). **J**. The number of cells preferring each target position in the “arrival” time window, among cells that have significant target position selectivity in that window (*p <* 0.01, permutation test for the target position GLM).

To investigate the temporal dynamics of the correlation across the trial, we computed the start-target tuning correlation between two halves of trials of the same neuron between one time window and another. Among neurons selective to both the start and the target position (*p <* 0.05 for both the start and the target GLMs, n = 174), the mean correlation between start position tuning early in the trial and target position tuning around target poke were positive (Figure 4E). For comparison, in the same group of neurons, the peak of the mean start-target tuning correlation among these neurons was 0.555, on the same scale as the target-target tuning correlation, which was 0.579, suggesting highly consistent start and target position tuning (Figure 4F&G).

The consistency of start position and target position tuning were not limited to strongly tuned neurons. Among all the neurons with spatial selectivity (*p <* 0.05 for any one of the GLMs, n = 808 ), the start tuning correlation, the target tuning correlation, and the start-target tuning correlation were all positively correlated (Figure S9). In other words, neurons tuned to the start position were more likely tuned to the target position, and the start and target tunings were more likely to be consistent. Collectively, these evidence suggested that single FOF neurons tracked the current head position.

One might notice the correlation between start position tuning late in the trial and target position tuning early in the trial (Figure 4E). This was due to the correlation between start and target positions in the task, which led to the weak ‘mirroring’ of the strong early-start to late-target correlation.

Start position encoding transitioned to target position coding mainly during the movement period (Figure 4H). We fit the neural spike counts across time to the start and the target GLMs, and defined the time of transition as the time when the *R*^2^ of the start position GLM first became smaller than the target position GLM (see Methods for details). For most of these neurons, the switch time was between the go sound and the target poke arrival time.

The numbers of neurons preferring each start position and target position spanned across all the ports (Figure 4I-J). The preferred positions were not uniformly distributed. There were more neurons with preferred start positions (*χ*^2^(6, *N* = 202) = 17.35, *p* = 0.009) and target positions (*χ*^2^(6, *N* = 209) = 48.39, *p* = 9.86 *×* 10^*−*9^) at the most leftward and rightward ports. Consistent with the current head position coding, the distribution of the preferred start position among start position selective neurons were similar to the distribution of preferred target position among target position selective neurons (*χ*^2^(6, *N* = 411) = 9.67, *p* = 0.139, Figure 4I&J).

### FOF neurons represented movement directions asymmetrically

Despite our ability to decode both the vertical and horizontal direction during planning (Figure 3B), single neurons in the FOF were not tuned equally to all movement directions (Figure 5A&B). First, there were significantly more neurons that preferred horizontal directions than those who preferred vertical directions (*χ*^2^(1, *N* = 274) = 76.91, *p* = 0 for the go window). Second, there were significantly more neurons who preferred downward directions than those who preferred upward directions (*χ*^2^(1, *N* = 274) = 5.37, *p* = 0.02 for the go window). Interestingly, rats mostly made errors into the same left/right side as instructed (Figure S2B) and to a lower direction (Figure S2C). For example, when instructed to top-left, errors were usually to the middle-left or bottom-left ports but rarely to the right. This type of behavioral asymmetry was consistently observed in other rats trained on similar tasks (Figure S3), and similar spatial asymmetries have been reported in human eye movements (Collewijn et al., 1988, Collewijn and Tamminga, 1984, Ke et al., 2013). Consistent with previous findings, there was no significant difference between the numbers of neurons preferring the ipsilateral and the contralateral side (*χ*^2^(1, *N* = 274) = 0.88, *p* = 0.34, Erlich et al., 2011).

**Figure 5.**
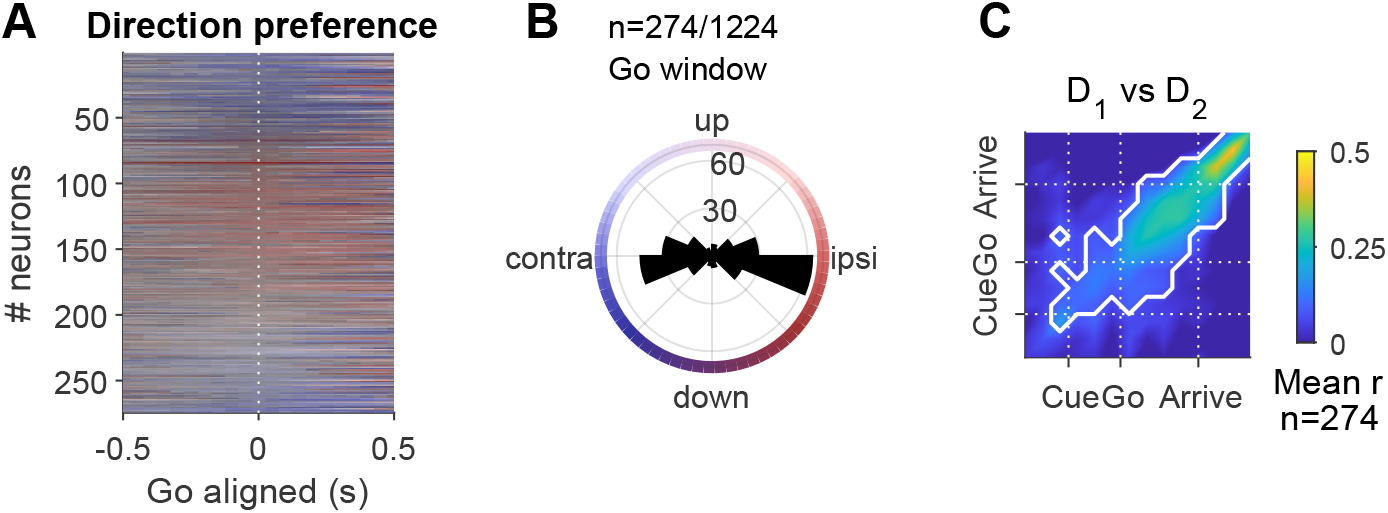
Direction preference of FOF neurons. **A**. The preferred direction of 274 cells that had significant direction selectivity in either the “post-cue” or “go” time windows (*p <* 0.01, permutation test for the GLM), in sliding windows aligned to the go sound (50 ms per bin, 300 ms bin size). The alignment was causal, i.e. time 0 indicated -300 to 0 ms. The color indicates the prefered angle (as in B.) and the saturation indicates the relative amplitude of the *R*^2^ of the direction GLM, and the neurons were sorted by the preferred direction in the “go” time window. **B**. The distribution of preferred direction in the “go” time window (0 to 300 ms after the go sound). **C**. Pearson correlation of direction tuning curves at one time versus another, among same neurons as in A and B. Color indicate the mean correlation across these neurons. White contour indicates the area with where correlation was significantly larger than zero with Bonferroni correction.

**Figure 6.**
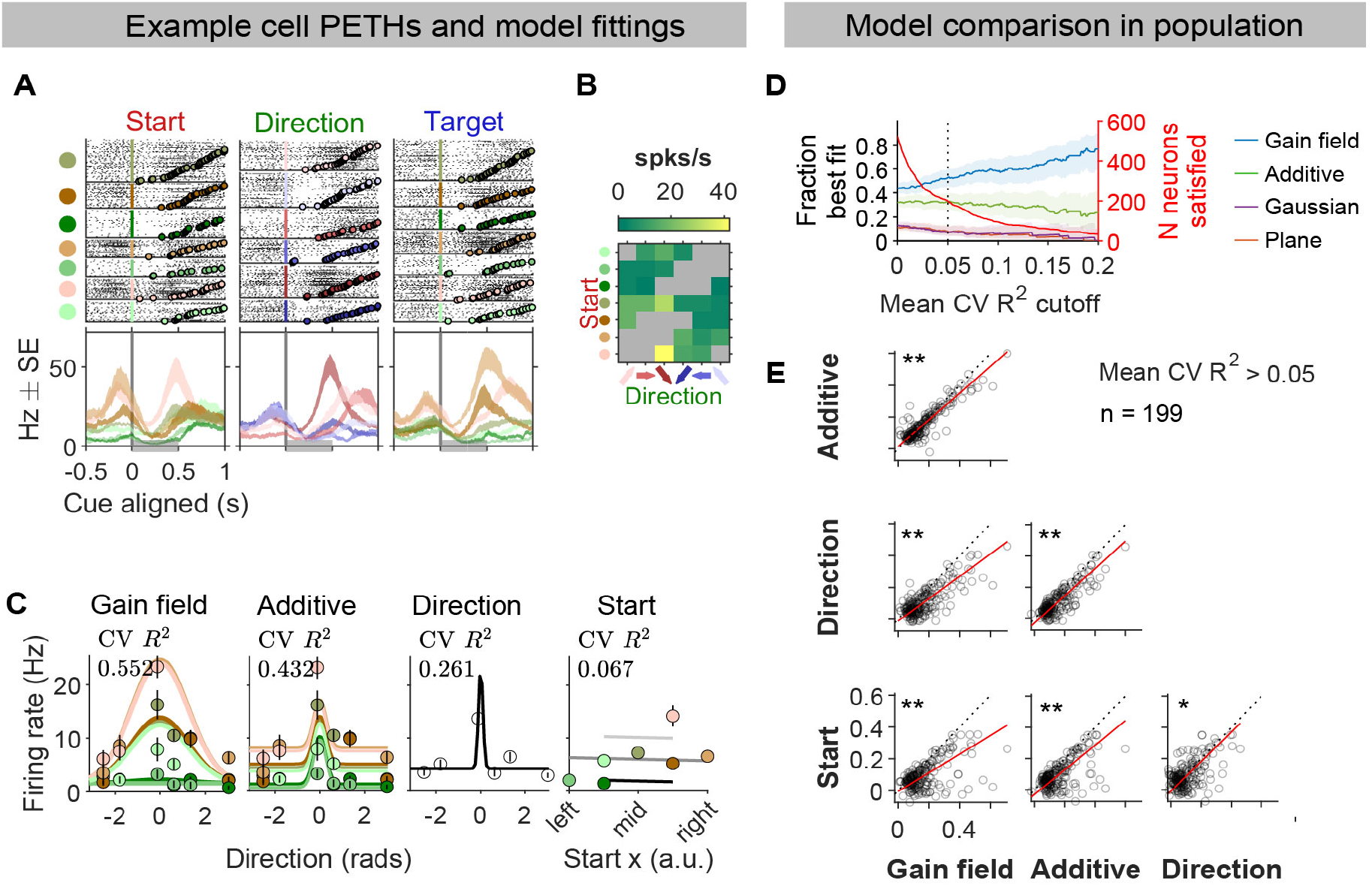
The spatial selectivity of FOF neurons were best explained by the gain field model. **A**. Raster plots and PETHs of an example neuron. Trials were grouped by the start position (left panel), the direction (middle panel) and the target position (right panel). Spikes were aligned to the visual cue onset. PETHs were generated by a causal half-Gaussian Kernel with an SD of 400 ms. The gray bar at the bottom indicates the 500 ms time window after the visual cue onset, the time window used for other panels in this figure. **B**. Estimated mean firing rate in each movement trajectory using a maximum a posterior estimator, same as in Figure 2. **C**. Predicted firing rates of four fit models (lines) and the mean and s.e.(circles and error bars) of firing rates in each trial condition. *CV R*^2^, cross-validated *R*^2^. **D**. Left axis, fraction of neurons best fit by the model, among neurons whose mean cross-validated *R*^2^s over the 4 models was larger than the x-axis indicated value. Best fit was defined as having the largest cross-validated *R*^2^ among the 4 models. Error bars were 95% confidence intervals of the binomial distribution. Right axis, neurons that crossed the mean *CV R*^2^ criteria for each x-axis value. **, *p <* 0.001. *, *p <* 0.05. **E**. Each panel plots the cross-validated *R*^2^s of the x-axis model versus the y-axis model. Each circle indicates a neuron. The red line indicates the total least square fit to the data. The dashed black line marks the diagonal. All single units were included in the analysis (n = 1224), but each panel only showed neurons whose mean of cross-validated *R*^2^s in the 4 models were larger than 0.05 (n = 199). Statistical test and total least square fitting were also based on these neurons. P values indicates the significance against the null hypothesis that the difference between the Fisher-z transformed cross-validated *R*^2^s in the x-axis model and the y-axis model was not significantly different from zero (permutation test of the mean). The mean *R*^2^ in the x-axis model was larger than the y-axis model in all of these panels.

### Mixed selectivity of movement direction and head position

In previous analyses, we examined the encoding and decoding of one spatial variable at a time, although single FOF neurons seemed to encode the conjunction of multiple spatial variables by visual inspection (Figure 2). Nonlinear mixed selectivity supports flexible readout by allowing high-dimensional representation of information from multiple sources (Rigotti et al., 2013). For example, a nonlinear transformation is required to compute distance 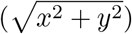 from neurons that encode the x and y position.

Gain field models are a common function form of nonlinear mixed coding of spatial information. They are so named since the information of one source multiplicatively modulates the information of another source (Andersen et al., 1990, 1985, Andersen and Mountcastle, 1983, Salinas and Sejnowski, 2001). For example, the primate frontal eye field encodes a gain field mixture of the initial eye-in-orbit position and the saccade vector (Cassanello and Ferrera, 2010, 2007). Following this, we hypothesized that FOF activity might be described as a gain-field mixture of the current position and the upcoming movement direction. We tested the hypothesis by comparing four encoding models: a pure direction tuning model (Eq. 1, see Methods), a pure position tuning model (Eq. 2), an additive mixed tuning model (Eq. 4), and a gain-field mixed tuning model (Eq. 3).

For each neuron, we fit spike counts in the 0 to 500 ms time window aligned to the visual cue onset to the four function forms (Figure 6A-C), and quantified the goodness of fit with the cross-validated *R*^2^ (*CV R*^2^). The vast majority of neurons were categorized as best fit by the gain field model, and the fraction of neurons best fit by a gain field model grew when the comparison was restricted to more spatially selective neurons (Figure 6D). Among neurons whose average *CV R*^2^s for the 4 models were larger than 0.05, the mean *CV R*^2^ for the gain field model was significantly larger than all the other models (Figure 6E, Table 1). The result was qualitatively the same for spike counts in the -300 to 500 ms window aligned to the go sound, as well as in the 500 ms window with maximum cross-trial-type variance aligned to the visual cue onset, demonstrating consistency across time. Thus, the majority of FOF neurons had nonlinear mixed selectivity to the self-centered and the world-centered spatial variables, consistent with findings in the primate frontal and parietal cortices during motor planning (Andersen et al., 1990, 1985, Andersen and Mountcastle, 1983, Salinas and Sejnowski, 2001).

**Table 1.** Difference of the Fisher-transformed *CV R*^2^s between model pairs for neurons whose mean *CV R*^2^ *>* 0.05 for all four models. Permutation test for the null hypothesis that *atanh*(*Model*1) *− atanh*(*Model*2) = 0 with 10^5^ permutations.

| $M_1$ | $M_2$ | $p$ value | $\text{atanh}(\text{Model1}) - \text{atanh}(\text{Model2})$ Mean, [95% CI] |
| --- | --- | --- | --- |
| Gain field | Additive | $2 \times 10^{-5}$ | 0.0129, [0.0073, 0.0229] |
| Gain field | Direction | $2 \times 10^{-5}$ | 0.0617, [0.0544, 0.0774] |
| Gain field | Start | $2 \times 10^{-5}$ | 0.0753, [0.0650, 0.0960] |
| Additive | Direction | $2 \times 10^{-5}$ | 0.0488, [0.0430, 0.0588] |
| Additive | Start | $2 \times 10^{-5}$ | 0.0624, [0.0529, 0.0775] |
| Direction | Start | 0.02 | 0.0136, [0.0025, 0.0259] |

### FOF population activity is most similar to a recurrent network with self-centered input and self-centered output

Gain fields have been suggested as a powerful computational mechanism for reference frame transformation, as it allows a downstream population to read out an arbitrary combination of the spatial information from different sources (Pouget and Sejnowski, 1997, Salinas and Abbott, 1995). That said, our current data doesn’t provide strong evidence whether the FOF is involved in the reference frame transformation or alternatively inherits representations from upstream regions. We turned to a computational modeling approach to gain insight into this question. We analyzed the activity of four types of networks (n=20 per class) trained on a similar task that as the visually-guided delayed-orienting task but with different input and output reference frame configurations (Figure 7A,S11): self-centered input and self-centered output (ego*→*ego); self-centered input and world-centered output (ego*→*allo); world-centered input and self-centered output (allo*→*ego); world-centered input and world-centered output (allo*→*allo). Thus, two of the network types needed to perform a reference frame transformation (ego*→*allo, allo*→*ego) and two of them did not (ego*→*ego, allo*→*allo).

**Figure 7.**
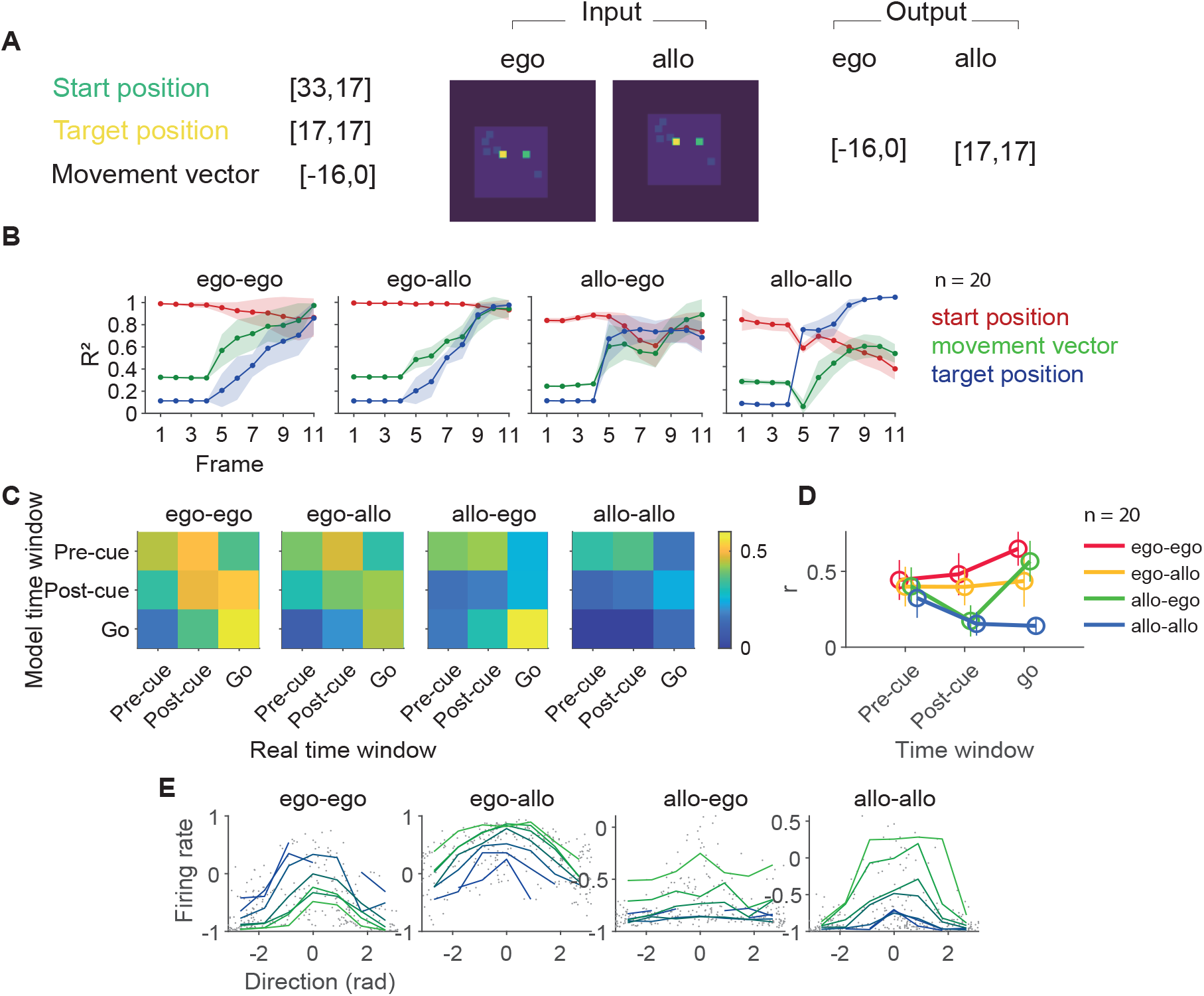
FOF population activity is most similar to a recurrent network with self-centered input and self-centered output. **A**. The input and output of an example trial in different networks. The start port (cyan) and the target port (yellow) coordinates are relative to the light-blue bounding box (the world frame). The movement vector is the displacement between the start port and the target port. The start port is always at the center of the image for ego-** networks, and the world frame is always at the center of the image for allo-** netowrks. **B**. The *R*^2^s between the decoded and the actual spatial variables. Shaded area denoted the s.e. over 20 training and testing epochs. The ego-ego network had the most similar temporal pattern of decoding accuracy as the FOF neurons. **C**. The representational similarity between the FOF neural population activity and the population activity in recurrent neural networks. The pre-cue, post-cue and go time windows in the networks were the 3rd, 7th and 11th time frames, respectively. The start position was visible throughout the 11 frames, and the target position (in the relevant reference frame) was visible briefly at the 4th frame. The representation similarity was computed with the the first 4 principle components of pseudo-population activity in the FOF neural data and the network hidden unit data. **D**. The representation similarity between the FOF neural population activity and the network hidden unit activity during corresponding time windows. Circles and error bars were mean and s.d. from 20 training and testing epochs. The means were the same as the diagonal elements in figure C. Representation similarity of the ego-ego models were significantly higher than other models in post-cue and go windows(Table 2). **E**. Example units in the four networks that had gain-field-like selectivity to start position and direction. Each dot is a trial and each line is the average over trials with the same start position which is indicated by color.

We decoded the start position, target position, and movement direction (the vector defined as target - start position) from the hidden unit activity during test trials in the same way as in real neurons, and found the ego-ego networks had the most similar temporal pattern of decoding accuracy as real neurons (Figure 7A). Representational similarity analysis (RSA) also showed the highest resemblance between real neurons and the hidden units of the ego-ego model (Figure 7B-D). RSA is a powerful technique to compare activity in networks (real or artificial) with different architecture, as long as the same stimuli can be presented to each network (Kriegeskorte, 2008). The idea behind RSA is to first estimate correlation between stimulus evoked activity patterns in each network and then to compare the patterns of correlations across the two networks. There was a significant effect of model on the representational similarity between the model and the real neural population (*F* (3, 236) = 41.6, *p* = 1.27 *×* 10^*−*21^, one-way ANOVA). The ego-ego model activity was significantly more similar to real neural population than all the other models in the post-cue window and the go window (Table 2, Welch’s t-test).

**Table 2.** Comparing the representational similarity for the ego-ego model versus the other three models. Welch’s t test for the alternative hypothesis that the mean of the representational similarity between the ego*→* ego model and real neurons were higher than those between another model and real neurons.

|  | pre-cue | post-cue | go |
| --- | --- | --- | --- |
| **ego-ego vs. ego-allo** | $t(38.00) = 1.05$<br>$p = 0.150$ | $t(37.34) = 2.02$<br>$p = 0.025$ | $t(32.95) = 4.69$<br>$p = 2.29 \times 10^{-5}$ |
| **ego-ego vs. allo-ego** | $t(37.90) = 1.02$<br>$p = 0.156$ | $t(35.03) = 7.96$<br>$p = 1.14 \times 10^{-9}$ | $t(36.79) = 2.11$<br>$p = 0.021$ |
| **ego-ego vs. allo-allo** | $t(38.00) = 2.86$<br>$p = 0.003$ | $t(29.77) = 9.19$<br>$p = 1.68 \times 10^{-10}$ | $t(26.86) = 18.41$<br>$p = 4.65 \times 10^{-17}$ |

Interestingly, all four models showed gain-field-like activity in some of the hidden units (Figure 7E), indicating that gain-field modulation is not uniquely associated with reference frame transformation within the network. These results suggested that the FOF does not transform preparatory activity from one reference frame to another, despite the observation of allocentric spatial encoding during the movement period.

## Discussion

Motor planning in rodents has previously been studied with tasks where the action space is either very high dimensional (such as navigating a maze or an open field) or very low dimension (such as 2AFC or go-nogo tasks). We took an intermediate approach with an orienting task that involved 6 movement directions and 7 head positions. We observed encoding of both egocentric and allocentric spatial parameters at the single-neuron and population level in the FOF. At the population level, two distinct 2D maps could be decoded from the neural activity in the FOF: a 2D map of current position on the poke wall and a 2D map of the future movement vector. The encoding of allocentric start position, egocentric movement direction and allocentric target position emerged sequentially over the trial. Despite the presence of allocentric information in the FOF, preparatory activity was in the egocentric reference frame.

This work presents a substantial advance in the conception of the function of the FOF. First, we established that preparatory activity in the FOF is encoded in a self-centered (egocentric) reference frame, consistent with the consensus view of preparatory activity in the primate frontal eye field (FEF; Schall, 2009). This preparatory activity is relatively abstract compared to the specific muscle sequence required for a movement: the upcoming direction of movement could be decoded ignoring the start position (Figure 3B). This finding is again consistent with findings from the FEF: auditory and visually guided saccades produce similar activity in the FEF even though they result in different eye/head kinematics (Russo and Bruce, 1994). Thus, although the preparatory activity is self-centered, neither the activity in FEF nor FOF are likely to represent a motor command, but instead represents an abstract command to shift attention to a part of space. Alternatively, it might not represent a command at all, but represent a utility or priority map which is used to generate movement commands in a downstream region, like the superior colliculus.

Our result builds on a recent interesting finding of world-centered spatial context modulating self-centered movement planning in rat M2 during navigation (overlapping anatomically with the region we define as FOF; Olson et al., 2020). The authors deduced that the M2 integrates spatial information toward the updating of planned movements.

However, their animals were not explicitly cued on each trial about the goal location or direction while navigating along a triple T-maze, which precluded them from demonstrating the temporal pattern of spatial encoding of different reference frames.

With a good experimental control over planning versus execution phases in our task, we demonstrated that planning is done preferentially in egocentric coordinates. Moreover, our comparison between the dynamics of the FOF neurons and the artificial neural networks using RSA casts doubt on the idea that the FOF transforms spatial information from one reference frame to another (Figure 3,7). A second difference is that we use a vertically oriented port wall instead of maze, which allowed us to demonstrate that the FOF contains two 2-dimensional maps (azimuth and elevation): one for position and one for direction (Figure 3B).

Although, it may appear that the FOF encodes the current position in a place-cell like fashion, there are ‘allocentric’ features of the box that could lead to this kind of tuning. For example, the right mystacial whiskers might touch the box edge when animals are in the right-most port or there could be visual cues that change (distance to the wall or box edges) depending on the position of the subject in the ports. That said, most of the position-tuned neurons had spatially smoothed tuning. For example, firing highest for the middle left port, but responding moderately to top left or bottom left (Figure 2K).

While it is not impossible for sensory responses to have this spatially smooth tuning (e.g. distance from an odour cue), it seems parsimonious to describe these as allocentric position tuned responses. Further experiments are required to better understand the nature of the allocentric tuning observed.

Subjects made few errors in the task, but when they did, they mostly made up/down errors, and three of the four rats made more errors downwards than upwards (Figure S2B&C, S3). At the neural level, there was an over-representation of left versus right than up versus down direction coding in the FOF, and an over-representation of downwards versus upwards directions (Figure 5). Although we do not have evidence for a causal relationship between the behavioral and neural observations, one might speculate that the left/right over-representation in the FOF might be behaviorally relevant (Jovalekic et al., 2011). In contrast to our findings, there were more pitch (up/down) tuned neurons than azimuth (left/right) tuned neurons reported in the M2 in rats foraging in a large arena (Mimica et al., 2018). There were two key differences between our experiment and theirs: first, they recorded from a larger anterior-posterior range of M2; second, our rats were under tight experimental control whereas their rats were foraging freely. Also, in our task, all visually-guided movements had a horizontal component which might have biased our results. It would be interesting to record the same neural populations in two tasks: one like ours and one like theirs to see if neurons in M2 can dynamically shift their tuning based on task demands.

The directional asymmetries in neural encoding and behavior suggest that a radially symmetric bump attractor model is insufficient as a computation model for the task (Wimmer et al., 2014). Plausible models for movement direction planning might be multiple discrete attractor models or ring attractor models that are asymmetric in connectivity weights: they might have stronger connectivity from upper direction preferring neurons to lower direction preferring neurons than the reverse, and have stronger connectivity between neurons that prefer the same left/right side than those preferring different left/right sides.

We found that allocentric and egocentric information is multiplexed as gain fields at the level of individual neurons. Gain modulation is a nonlinear process in which neurons combine information from two or more sources. Gain fields have been observed in a plethora of primate cortical and subcortical brain regions (for review, see Salinas and Sejnowski, 2001), including the primate FEF (Cassanello and Ferrera, 2010, 2007). While conjuctive coding of allo- and ego-centric information has been found in rodent para- and post-subiculum and the medial entorhinal cortices (Gofman et al., 2019, Peyrache et al., 2017) they were not explicitly modeled as gain fields even though theoretical work suggests that they might take that form (Bicanski and Burgess, 2020). According to the theory of gain field modulation, a neural population downstream of a population with gain field encoding could easily compute an arbitrary combination of the two spatial variables (Alexander et al., 2023, Pouget and Sejnowski, 1997, Salinas and Abbott, 1995, Zipser and Andersen, 1988). However, our comparison of recurrent networks showed that networks that do not perform reference frame transformation could also have gain fields (Figure 7).

Our task was inspired by paradigms widely used in non-human primates to study the neural mechanism of saccadic eye movements (Bruce and Goldberg, 1985) and rodent head orienting is, like a saccade a shift in overt attention, in that it redirects the sensory fields of vision, audition, olfaction and whiskers (Bush et al., 2016, McCluskey and Cullen, 2007, Monteon et al., 2010). Based on similarities in connectivity and function, the rodent FOF is a proposed functional analogue to the primate FEF (Ebbesen et al., 2018, Erlich et al., 2011, Reep et al., 1987, 1990), although a strict correspondence between rodent and primate frontal regions may not exist (Barthas and Kwan, 2017, Wise, 2008). Our findings of gain field modulation of movement direction by initial head position support the analogy, as this is similar to the observation that eye-in-orbit position gain modulates the planned saccade vector in FEF (Cassanello and Ferrera, 2010, 2007). One may argue that for the FEF both the initial gaze direction (or eye-in-orbit position) and the saccade vector are egocentric, and most of the reports about spatial encoding in the FEF was egocentric (but see Bharmauria et al., 2020). However, body positions in primate experiments are typically restricted by a primate chair. Recent work in freely moving monkeys found widespread world-centered coding across frontal and prefrontal cortex (Maisson et al., 2022), including supplementary motor area and dorsolateral prefrontal cortex. Although FEF was not one of the regions they recorded from, the finding suggests that allocentric representations are common across the brain, when animals’ movements are not restricted. Thus, we might speculate that FEF saccade planning activity in monkeys freely moving in an environment could be modulated by the current position of subject in the environment.

The observations of gain fields in the FOF, as well as the sequential encoding of movement direction and target position, were largely consistent with observations in primate motor planning more generally (Andersen et al., 1985, Bharmauria et al., 2020, Caruso et al., 2018, Salinas and Sejnowski, 2001, Wang et al., 2007, Zipser and Andersen, 1988). However, despite decades of research, a full understanding of phenomena like gain fields how they could causally contribute to reference frame shifts has been hampered by a lack of tools for precise perturbations for circuit dissection in primates (as discussed in Shenoy et al., 2013). In contrast, the neural circuits underlying basic phenomena of motor planning (i.e. spatial memory and movement initiation in 2AFC tasks), have been revealed in rodents models (Bharmauria et al., 2020, Duan et al., 2021, Guo et al., 2017, Inagaki et al., 2019, Kopec et al., 2015, Li et al., 2016, Yang, 2022). Our behavioral paradigm and neurophysiological observations provide a basis for employing the latest tools of rodent systems neuroscience to understand complex and ethological motor planning.

## Materials and Methods

### Subjects

Three adult male Sprague-Dawley rats and one adult male Brown Norway rat (Vital River, Beijing, China) was used in this study. For a portion of the experiments presented here, rats were placed on a controlled-water schedule and had access to free water 20 minutes each day in addition to the water they earned in the task. For some experiments, rats were given free access to a 4% citric acid solution (Reinagel, 2018), in addition to the normal water they earned in the task. They were kept on a reversed 12 hour light–dark cycle and were trained during their dark cycle. Animal use procedures were approved by New York University Shanghai International Animal Care and Use Committee following both US and Chinese regulations.

### Behavior

Rats were trained in custom behavioral chambers, located inside sound- and light-attenuated boxes. Each chamber (23 *×* 23 *×* 23 cm) was fit with a vertical 2-D port wall that had 7 operant ports and 1 reward delivery port, with speakers located on the left and right side (Figure 1). Each operant port contained a pair of blue and a pair of yellow light emitting diodes (LED) as visual cues, as well as an infrared LEDs and photo-transistors for detecting rats’ interactions with the ports. The reward delivery port contained a stainless steel tube for delivering water rewards.

The task timeline is described in detail in Results and Figure 1. In one rat (2147), in addition to the main type of timeline, which we denote as “target during fixation”, there were two other types of trial timelines: “target before fixation” and “target after go”. These trial types are described in detail in Figure S5.

The duration of the fixation period was dynamically adjusted for each animal, ranging between 0 *∼* 1.2 s (Figure S2). A trial was considered a fixation violation if the rat withdrew from the start port before the go sound. In fixation violation trials, an “error” sound was delivered and the trial was aborted.

In the final behavioral stage, three rats (subject ID 2068, 2095, 2134) moved in 6 directions and 30 movement trajectories, and one rat (subject ID 2147) moved in 4 directions and 16 movement trajectories (Figure S1).

In rat 2068, 2095 and 2134, there were two session types interleaved across days: the “reference” sessions and the “distance” sessions. In the “reference’’ sessions, each of the 6 directions had 4 movement trajectories of the same distance. In the “distance” sessions, each of the 6 directions had 3 movement trajectories involving 3 ports, where the distance of one movement trajectory is twice of the movement of the other two(Figure S1).

In rat 2147, there were 4 movement directions, from and to 7 operant ports. Each direction had 4 trajectories of the same distance, summing up to a total of 16 trajectories (Figure S1).

### Behavior training pipeline

Rats went through a series of training stages, which had mainly two phases: (1) the operant conditioning phase, and then (2) the multi-directional orienting phase.

In the operant conditioning phase, rats became familiar with the training apparatus and learned to do a one step poke into the illuminated choice port. The first stage was to learn to collect reward from the reward delivery port. Each trial began with the illumination of the reward port, and water reward was immediately delivered upon port entry, followed by a short timeout period before the start of next trial. After the rats learned to poke into the reward port reliably (not missing any reward for 6 trials in a row), they proceeded to the next training stage. In the second stage, we turned on the LED for several random ports at the beginning of each trial. Rats had to first poke into any illuminated choice port before gaining water reward from the reward delivery port. The number of the illuminated port will gradually decrease to one after several trials when animals started to learn. After animals were able to poke the only illuminated choice port successfully for 6 trials in a row, we will upgrade them to the second training phase.

In the first stage of the multi-directional orienting phase, the start port was always the central port, and the target port was one of the 6 surrounding ports. Rats needed to poke into the start port to trigger the target port light, and then poke into the target port after a delay. Trials of the same movement trajectory was repeated until the animal could do several correct trials in a row. The training of “fixation” at the start port was introduced in this phase. Fixation means the rat had to keep its nose in the start port for a given time period (typically *>* 0.5*s*). Fixation duration was initially 0.02 s early in the training, and was gradually increased based on an adaptive steps method: the fixation duration would increase on a successful fixation, and decrease when the fixation failed. We trained subjects to perform fixation for at least 0.6 s before the surgery, and fixation duration always jittered across trials in recording sessions. However, the speed to recover fixation after the surgery varied across subjects, thus we manually adjusted the fixation duration for each subject. In 2095, the mean fixation period in each session was shorter than 0.2 s for around 30% of sessions. In other subjects, the mean fixation period was typically longer than 0.4 s (Figure S2 G). In the second stage of the training phase, the start port could be any one of the 7 ports, and the target port was one of the 6 remaining ports. Rats were trained on the “target during fixation” trial class. Rat 2147 was then introduced to the “target before fixation” and “target after go” trial classes, described in Figure S5.

### Electrophysiology

Rats were unilaterally implanted with Cambridge Neurotech microdrives and 64 channel silicon probes. To target the frontal orienting field (FOF), silicon probes were placed at anteroposterior (AP) and mediolateral (ML) coordinates as following: rat 2068, AP +2.2, ML +1.5; rat 2095, AP +2.0, ML +1.5; rat 2134, AP +2.5, ML -1.5; rat 2147, AP +2.2, ML -1.4 (Figure S4). The probes were placed initially 0.5 mm below the brain surface and were advanced 50 to 100 um every 1 to 4 days, after each recording session. The same neurons could be sampled across sessions.

### Analysis of neural data

#### Spike sorting

Spike sorting was performed with automatic clustering and manual curation in JRClust. Single units were defined as those with signal-to-noise ratio larger than 5, fraction of inter-spike interval smaller than 1 ms less than 1%, and within-trial firing rate larger than 1 Hz. However, our main results were robust to different single neuron criteria (Figure S6).

#### Data inclusion

For a unit to be included in the main figures, the session must have at least 8 trials for each of the 7 start positions, 4 or 6 movement directions and 7 target positions. The sessions could be “reference”, “distance”, or sessions from rat 2147. As described above, there were 3 types of trial timeline in rat 2147, but only “target during fixation” trials are included in the main figures. This resulted in 104 sessions and 1224 single cells in 4 animals.

In pseudopopulation decoding, we included cells from sessions that had at least 8 trials for each of the 6 movement directions, 7 start positions, and 7 target positions. The sessions could be “reference” or “distance” sessions. This resulted in 1197 cells from 99 sessions, 3 animals.

Video tracking was available in 58 sessions in 3 rats.

For neuron inclusion criteria for all the main figures, see Table S1.

Unless otherwise specified, only correctly performed trials were included in the analysis.

### Key time windows

In single-neuron level analysis for single spatial variables, we focused on 4 key time windows: *pre-cue*, -300 to 0 ms aligned to target cue onset; *post-cue*, 0 to 300 ms aligned to target cue onset; *go*, 0 to 300 ms aligned to the go sound; *arrival*, -150 to 150 ms aligned to arrival at the target.

When comparing the pure and mixed selective models (Figure 6), we used the 0 to 500 ms time window aligned to the visual cue onset in.

For continuous time windows, neural response was quantified by counting the number of spikes in sliding windows of 300 ms width and 50 ms step, aligned causally to task events, unless otherwise specified. Causal alignment means that the value at time 0 refers to the neural activity in a time bin between -300 ms and 0 ms. The responses could be aligned to the time of visual target cue onset, the go sound onset, the time of “fixation out” (the time when the nose left the start port, detected by the IR sensors) or target poke (the time when the nose arrived at the target port, detected by the IR sensors).

### Cross-validated *R*^2^s and log likelihood

The cross-validated *R*^2^s (denoted as *CV R*^2^s in figures) was defined as

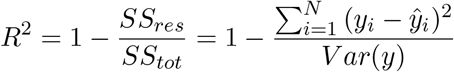

*ŷ*_*i*_ was the predicted mean spike count in the *i*th trial from the test set, and *y*_*i*_ was the observed spike count in this trial. In the best case, *y* is equal to *ŷ*, and the cross-validated *R*^2^ is 1. In the worst case, *ŷ* is uncorrelated with *y*, and 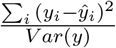 can be larger than 1 by chance, thus *CV R*^2^s can be negative.

The cross-validated log likelihood is defined as

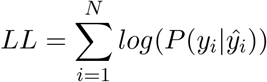

*P* (*y*_*i*_|*ŷ*_*i*_) denote the probability of observing the spike count on the *i*th trial being *y*_*i*_, given the predicted mean spike count being *ŷ*_*i*_. *N* denote the number of trials. We assumed spike counts to follow the Poisson distribution.

### PETHs

For each neuron, we combined the spike trains in correctly performed trials by the movement directions or target positions, and generated smoothed PETHs with an half-Gaussian kernel of 200 ms standard deviation. The kernel was causal, such that selectivity at time *t* was contributed by neural activity at or before time *t*.

### Generalized linear models for single neuron selectivity

We fit the neural spike counts in specific time windows to 3 generalized linear models ( GLMs) with Poisson distributions:

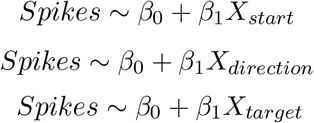

The spatial variables were all included as dummy variables. Start position had 7 levels, direction had 6 levels, and target position had 7 levels.

In the main text, we labeled neurons as “having significant selectivity to a spatial variable”. This significance i s derived from the p ermutation test of the GLM, where the test statistic was the leave-one-out cross-validated log-likelihood, against the null hypothesis that the log-likelihood was not significantly different from when trial labels were shuffled. For each spatial variable, the trial labels were shuffled 1000 times to obtain a distribution of goodness-of-fits, then the p value of the GLM was the fraction of log-likelihoods from shuffling that is greater than the log-likelihood from actual data.

The best selectivity of a neural response (Figure 3A) was assigned to the spatial variable with the smallest p value of the GLM. When there were ties in *p* values, we additionally compared the leave-one-out cross-validated log-likelihoods, and the best selectivity was assigned to the spatial variable with the largest sum of log-likelihoods. The reason for possible ties is that we are using permutation tests with 5000 permutations.

In Figure 3 B, the *R*^2^s were derived from GLMs fit similarly, but without cross-validation. Only the model with the largest *R*^2^ was plotted with the corresponding color at each time point. Both the best GLM measurement and the *R*^2^ measurement for the relative strength of spatial selectivity were robust to the task-induced correlation between spatial variables, which was verified with surrogate data (Figure S7, also see Surrogate data paragraph below).

Cross-validated *R*^2^s of the GLMs in Figure 2 were derived from 10-fold cross validation.

### Spatial preference of single neurons

The preferred direction of a neuron in a specific time window was defined as the direction of the weighted vector sum of the coordinates of the 6 movement directions, where the weight for each direction was the mean firing rate in that direction. Horizontal directions were defined as directions between *−*45° and 45° around the horizontal directions, and so was for vertical directions. Downward directions were directions that were *−*90^*◦*^ to 90^*◦*^ around the downward direction, and so was for upward directions. The preferred position of a neuron at a specific time window was simply denoted as the port with the highest mean firing rate.

### Start-target tuning correlation

The start-target tuning correlation in Figure 4D was defined as the Pearson correlation between the tuning curve for start position in the “pre-cue” window and target position in the “arrival” window:

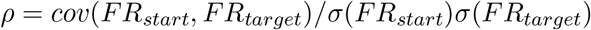

*FR*_*start*_ is a vector where each element is the mean firing rate for a specific start position, and *FR*_*target*_ is a vector where each element is the mean firing rate for a specific target position. Similarly, we calculated the start-target tuning correlations for different pairs of time windows and the tuning correlation between time *t*_*i*_ and *t*_*j*_ for the same variable. The time windows were warped to align to the time of the visual cue, the go sound and the target poke. The 300 ms “pre-cue” and “arrival” time windows were preserved. For these time windows, we calculated the tuning correlations between to halves of trials that were randomly split, that is, we used start tuning in the first half versus target tuning in the second half, and vice versa, and then took the average of the two correlation coefficients.

The tuning correlation was subject to the Fisher-z transformation, and then tested against the null hypothesis that the mean value across a specific neural population was not significantly larger than zero. The p value indicated the fraction of mean lying below zero among 10^4^ bootstraps, and was corrected with Bonforroni correction.

### Timing of start-target switching

In Figure 4 I, spike counts in 300 ms time windows of 50 ms steps were fit to the Poisson GLM of either start or target position. The *R*^2^s of each GLM across time was smoothed with a moving average kernel with the size of 3 bins, and then the start GLM *R*^2^ was subtracted by the target GLM *R*^2^. The time of switching from start encoding to target encoding was defined as the first time window where 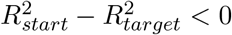, after the positive peak of 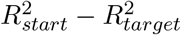.

### Mixed selectivity

To detect mixed selectivity for each neuron, we compared between 4 models of the neural firing rate.

In the Gaussian direction tuning model, the firing rate was a Gaussian function centered by the preferred direction, defined as in the “Spatial preference of single neurons” section in Methods.

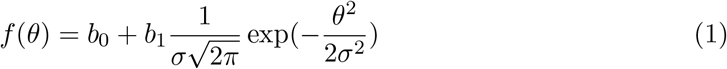

In the start position plane model, the firing rate was modulated linearly by the horizontal and vertical coordinates of the start position.

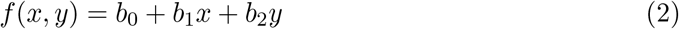

In the gain field model, the firing rate was a multiplicative combination of a Gaussian tuning centered by the preferred direction and a start position modulation.

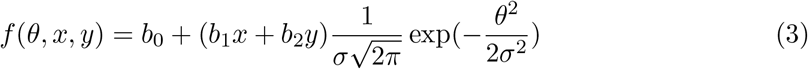

In the additive model, the firing rate was an additive combination of a Gaussian tuning centered by the preferred direction and a start position modulation.

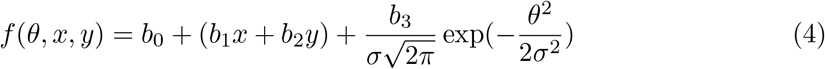

In these functions, *f* was the firing rate, *θ* was the movement direction relative to the preferred direction, and *x* and *y* are the horizontal and vertical coordinates of the start positions in the port wall. *b*_0,1,2,3_ and *σ* were fit for each model with maximum likelihood estimation assuming Poisson spike counts, using the Matlab *fmincon* function. The models were fit with 20-fold cross validation.

For each pair of models, we compared the cross-validated *R*^2^s among neurons whose average of cross-validated *R*^2^s of the 4 compared models was larger than 0.05. Significance level was tested against the null hypothesis that the mean difference between the Fisher-z transformed cross-validated *R*^2^s in the x-axis model and the y-axis model was not significantly larger than zero, quantified as the fraction of the mean value smaller than zero among 10^5^ bootstraps.

For a given movement direction, the target position coordinates were simply an additive translation of the start position, thus the start position and target position modulations were the same in the gain field model and the additive model.

### Pseudopopulation decoding

Pseudopopulations were constructed separately when decoding each spatial variable. For example, when decoding the start position, we constructed the pseudopopulation by resampling trials for each start position. For each neuron, trials were randomly split into 2 folds for each condition of the spatial variable, and 32 trials were resampled for each fold in each condition, yielding 224 trials per fold for position decoding and 192 trials per fold for direction decoding. These resampled trials were termed “pseudo-trials”. Pseudo-trials were resampled randomly with a different seed for each neuron, so as to remove trial-by-trial correlations between neurons in the same session.

We included single neurons from sessions that had at least 8 trials for each of the 7 start positions, 6 directions and 7 target positions (1197 neurons, 99 sessions, 3 rats). In each pseudopopulation, neurons were also resampled with a different seed, so that only about 63.2% of the neurons were included in each pseudopopulation. We generated 100 pseudopopulations for the decoding each spatial variable.

The error of decoding was defined as the Euclidean distance between true and predicted coordinates. The goodness of prediction was measured as the mean error between the predicted and the actual coordinates across all the pseudo-trials in that pseudo session. For start and target position, we removed the trials with [0 0] coordinates from the quantification of decoding accuracy. This is because predictions from unsuccessful decoding tend to cluster at the [0 0] coordinates, so decoding accuracy for the [0 0] coordinates in positions (i.e. the central port) will always seem “good”.

Decoding was performed with multivariate regression models (*mvregress* function in Matlab Statistics and Machine Learning Toolbox). The spike counts of the training set and the test set were first combined, z-scored, and applied to principle component analysis, and then split again for training and decoding. We found that the cross-validated decoding accuracy was the best when including only the first 4 PCs when decoding start position from neural activity in the “pre-cue” time window (Figure S8G), so we used 4 PCs for all subsequent pseudopopulation decoding.

In Figure 3B, multivariate regression models were trained and tested with spike counts in 3 key time window as indicated in the plot (see “Key time windows” section for details). The plots showed the result from one example pseudo-session, the same pseudo-session as the first row in Figure 3C. In Figure 3C, the time windows were 300 ms width with 50 ms step, causally aligned to the go sound. Each row was a different pseudo-session. Each row of the heat map was a pseudopopulation with a different resampling of neurons, and the color indicated the mean error with two-fold cross-validation. In Figure 3D, the mean error of one spatial variable was subtracted by the mean error of another spatial variable in each pseudopopulation, termed the Delta error, which represented the relative goodness of decoding between the two spatial variables. The distribution of Delta errors of the 100 pseudopopulations was then illustrated with the mean and standard deviation.

In Figure 3E, models were trained at one time window and tested at another also with two-fold cross validation. In the last panel, a multivariate regression model was trained to decode start position coordinates from the start position pseudopopulation data, and then tested its decoding of target position coordinates from the target position pseudopopulation data. The heat-map showed the average of mean errors across 100 pseudopopulations. P values for multiple comparisons were obtained with extreme pixel-based permutation test (Cohen, 2014). One dummy heat-map was generated for each pseudopopulation by shuffling trial labels, and the maximum of each heat map was gathered to construct a null distribution. The white contours enclosed the area in which the averaged mean error was smaller than the minimum value of this distribution (*p <* 0.01).

### Video tracking

Videos were acquired at 30 frames per second with one Raspberry Pi Camera at the top of the rig.

We estimated the coordinates of the head, the ears and the hip of the rat in the video frames using DeepLabCut (Mathis et al., 2018). Coordinates were in the unit of pixels. The head position was approximated as the Intan chip plugged on the animal’s head. For continuous sampling of body coordinates aligned to task events, we linearly interpolated to achieve the sampling rate of 100 frames per second.

### RNN

We trained recurrent neural networks with 100 hidden units using pytorch (nn.RNN). The input to the networks (at each time) was a 100-by-100 pixels image downsampled to 40-by-40 pixels. The hidden layer was connected to a two node linear output layer, representing coordinates (x,y).

In the training set, the start port and target port coordinates were randomly generated inside a “port wall” bounding box, which was also the world frame. The image itself was the visual frame. The input image, which was the visual frame, had a dark background with an inner frame denoting the world frame. The start port, target port, and the distractors were denoted as intensities. Specifically, the start port had an intensity of 0.7, the target port had an intensity of 1, and the distractor had an intensity of 0.2. These elements were plotted as pseudo-colors in the figures. In the test set, the start port and target port corresponded to the 24 trial types in the “reference” sessions.

The networks were trained to take flattened image inputs over 11 time frames and to report the 2-D spatial coordinates at the last time frame (Figure 7A,S11). We presented the start port on the first frame, which stayed on for all 11 frames, and the target port was presented only on frame 4. The nonlinearity used in the model was tanh, and stochastic gradient descent was employed during training.

### Reference frames of the models

There were 4 task configurations that differed in the reference frame of inputs and outputs, namely: self-centered input, self-centered output (ego-ego); self-centered input, world-centered output (ego-allo); world-centered input, self-centered output (allo-ego); world-centered input, world-centered output (allo-allo).

For the ego-input networks, the start port was always centered to the image (i.e. the visual frame), thus the target coordinates were the same as the movement vector. For the allo-input networks, the world frame was always central to the image, thus the world-centered target port position is directly readable from its position in the visual frame.

### RSA on RNN data

To examine the representation similarity between the hidden units in the RNN and the real neurons, we focused on the hidden unit activity at frame 3, 7 and 11. These frames corresponded to real neural activity during the pre-cue, post-cue and go time windows in the task, respectively.

We first computed the first 4 principle components of the hidden unit activity and real neuron activity separately, under the 24 trial types. We then constructed the representation dissimilarity matrix (RDM) for the two datasets by calculating the correlation coefficients for each pair of trial configurations. Finally, we computed the Pearson correlation coefficient for the linearized lower-triangular matrix for the two RDMs, as the measurement for the similarity between neural representation of the RNN and the real neurons.

### Surrogate data

To verify the reliability of our model comparison approaches, we performed the same analysis on surrogate data which were known to have specific spatial tuning as on real neurons. These surrogate spike counts were generated to have specific spatial tuning, to match the overall firing rate of a real neuron, and as if sampled from a real session.

To generate surrogate data matching one neuron but tuned to a specific model, we first fit the spike counts of the real neuron to that model, so as to obtain the predicted firing rate at each trial condition. We then randomly selected a real session and used a Poisson random process to generate spike counts that followed these predicted firing rates. In the surrogate data for GLMs, we only used neurons that have significant selectivity to that GLM to generate surrogate data. In surrogate data to compare between pure and mixed selectivity models, we used all neurons to generate surrogate data.

Following stages of analyses was identical for surrogate data and real data. If our methods were reliable, the surrogate data designed to have a specific functional form would be best fit by the same functional form.

## Code and data sharing

Please visit https://github.com/erlichlab/fof-visually-guided to access the code used for analyses and to generate figures. Links to the data are available from the github repository.

## Author Contributions

JCE and LL conceptualized and designed the experiments. LL, CM, JCE and JL collected the experimental data. LL, CM and JL curated the spike sorting and contributed to analysis pipelines for electrophysiology. LL analysed the behavioral and neural data under supervision of JCE. CM extracted tracking information from videos. HC developed an earlier version of this task. YC contributed to an earlier version of the spike sorting pipeline. TF did the RNN modeling with input from JCE and LL. LL and JCE wrote the manuscript with comments from all authors.

## Acknowledgements

The authors thank Yingkun Li, Nengneng Gao and Anyu Fang for their assistance with behavioral training and animal care. We thank Xuan Wen, Cequn Wang, Yidi Chen, Josh Moller-Mara, and Sylvain Dubroqua for their help with developing hardware and software for the training facility. LLJL and JL were supported by the NYU Shanghai Doctoral Fellowship, Winston Foundation Fund, and BOCO Fund for Science and Research. The research was also supported by grants to JCE from: the Program of Shanghai Academic/Technology Research Leader (15XD1503000); the Science and Technology Commission of Shanghai Municipality (15JC1400104); the 111 Project, Base B16018; the National Natural Science Foundation of China (NSFC; 31970962); NYU-ECNU Institute of Brain and Cognitive Science at NYU Shanghai. JCE also acknowledges the funders of the Sainsbury Wellcome Centre (The Wellcome Trust and the Gatsby Charitable Foundation).

## Supplemental Figures

**Table S1.** Criteria for the inclusion of neurons in each analysis. Single neurons: SNR*>*5, in-trial firing rate *>* 1 Hz, fraction of inter-spike interval less than 2 ms *<* 1%. All the cells are from sessions where the target cue illuminated during the fixation period.

| Figure | Panel(s) | N | Criteria |
| --- | --- | --- | --- |
| FIGURE 3 | A | 1224 | Single neurons in 4 animals that had at least 8 trials for each condition. 771 neurons from reference sessions, 426 from distance sessions, 27 from rat 2147. |
| FIGURE 3 | B-E | 1197 | Single neurons that had at all 6 directions, and at least 8 trials in each group of trial conditions. 771 neurons from reference sessions, 426 from distance sessions. 3 rats. |
| FIGURE 4 | D (black bars)-H | 174 | Single neurons that have significant selectivity to start port during the “precue” time window and target port during the “arrival” time window (GLM, both $p < 0.05$), among the 1224 neurons in FIGURE 3A. |
| FIGURE 4 | D (white bars) | 289 | Single neurons that have significant selectivity to either the start port during the “precue” time window or the target port during the “arrival” time window (GLM, only one $p < 0.01$), among the 1224 neurons in FIGURE 3A. |
| FIGURE 4 | I | 202 | Single neurons that have significant selectivity to start port during the “pre-cue” time window (GLM, $p < 0.01$), among the 1224 neurons in FIGURE 3A. |
| FIGURE 4 | J | 209 | Single neurons that have significant selectivity to target port during the “arrival” time window (GLM, $p < 0.01$), among the 1224 neurons in FIGURE 3A. |
| FIGURE 5 | A-C | 274 | Single neurons that have significant selectivity to direction during the “post-cue” or “go” time window (GLM, $p < 0.01$), among the 1224 neurons in FIGURE 3A. |
| FIGURE 6 | E | 199 | Single neurons that have the average of $CVR^2$s larger than 0.05. Effectively from 4 rats. |

**Figure S1.**
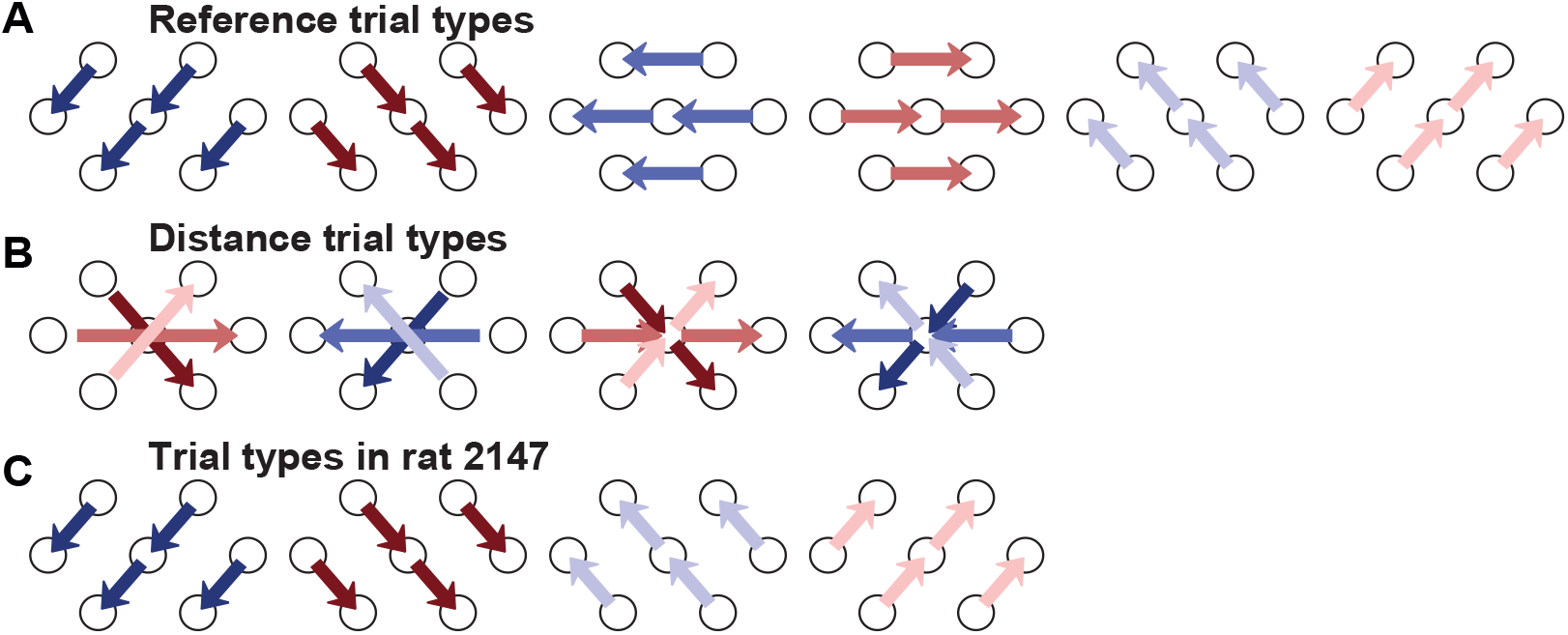
Movement trajectories configurations in each type of sessions. There were 24 configurations in the “reference” sessions. There were 18 configurations in the “distance” sessions, where each movement direction has two short distance configurations and one long distance configuration. In subject 2147, there were 16 configurations.

**Table S2.**
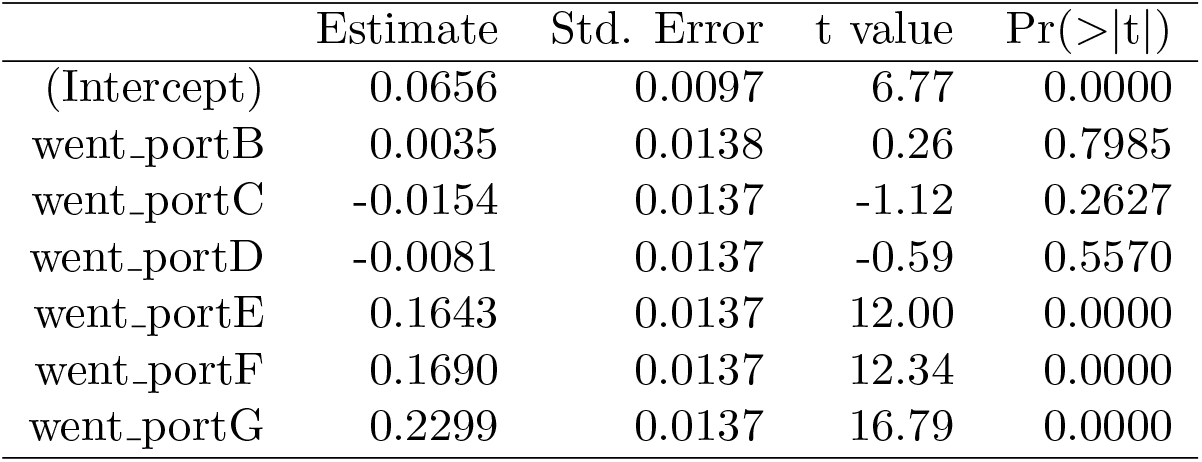
Fraction of poking into each port quantified with the linear model: *fraction ∼ went*.*port*. Each observation was the fraction of errors into a specific port for one subject. Data from all the error trials in the 31 “rand” group subjects was included.

|  | Estimate | Std. Error | t value | Pr(>\|t\|) |
| --- | --- | --- | --- | --- |
| (Intercept) | 0.0656 | 0.0097 | 6.77 | 0.0000 |
| went_portB | 0.0035 | 0.0138 | 0.26 | 0.7985 |
| went_portC | -0.0154 | 0.0137 | -1.12 | 0.2627 |
| went_portD | -0.0081 | 0.0137 | -0.59 | 0.5570 |
| went_portE | 0.1643 | 0.0137 | 12.00 | 0.0000 |
| went_portF | 0.1690 | 0.0137 | 12.34 | 0.0000 |
| went_portG | 0.2299 | 0.0137 | 16.79 | 0.0000 |

**Figure S2.**
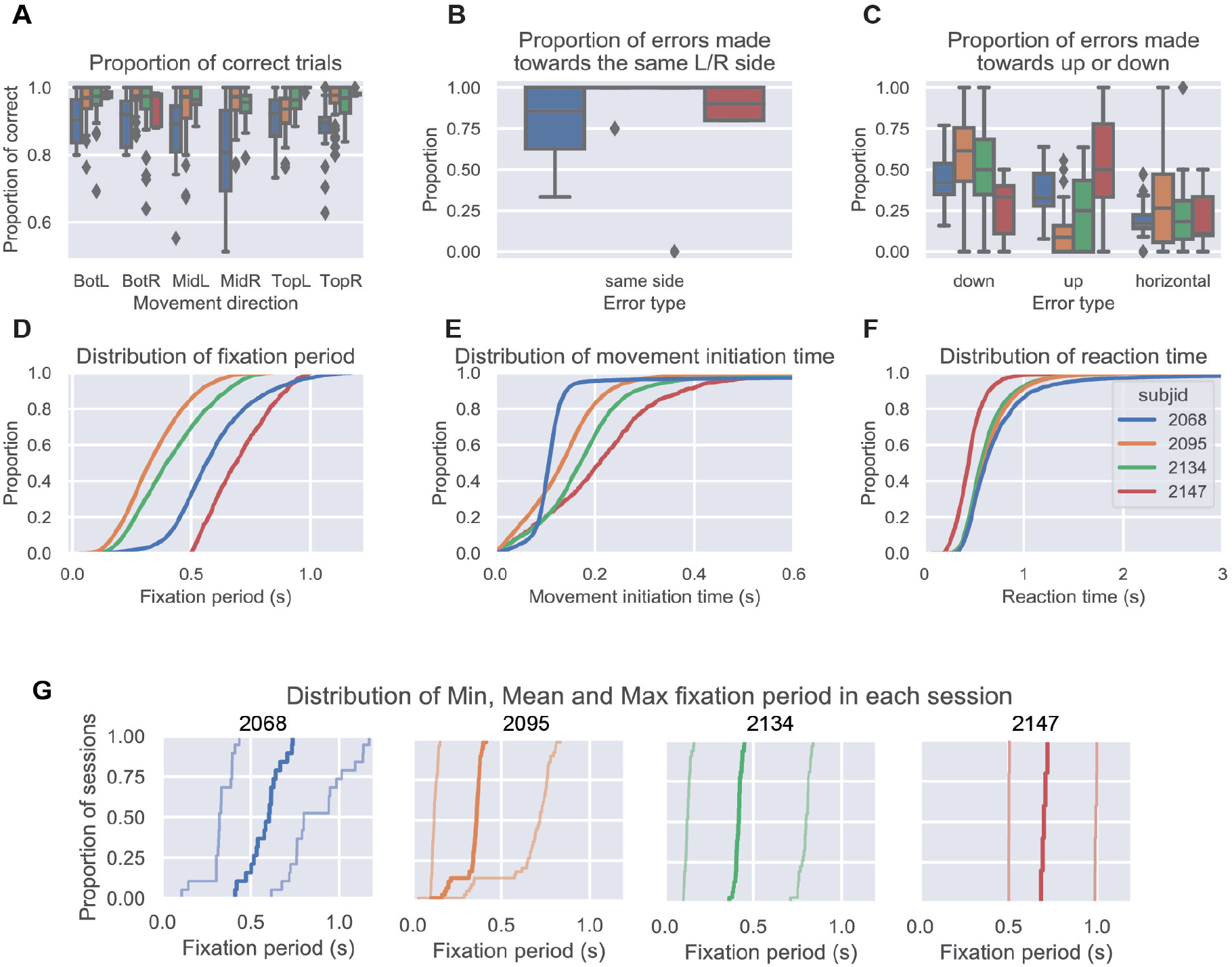
Behavioral measurements of the 4 animals. **A**. The proportion of correct trials among all the finished trials in each movement direction and each animal. The proportion of correct movements among all finished trials in each session was summarized by subjects. **B**. The proportion of errors made into the same side as instructed, among trials starting from the central port. Percentage of error trials into the same side was 93.55% ± 0.17% (*mean ± s.d*.) across the 104 sessions. **C**. The proportion of errors that were relatively up, down or in the same vertical level, compared to the correct target port position among all error trials. In A-C, points indicate outliers and box plot indicates the quartiles. **D**. The distribution of instructed fixation period in correctly performed trials, defined as the time between nose arrival at the start port and the go sound. **E**. The distribution of movement initiation time, defined as the time between the go sound and the nose departure from the start port. **F**. The distribution of reaction time, defined as the time between the go sound and the nose arrival at the target port. The nose departure or arrival time were detected by the infra-red sensors in each port. **G**. The distribution minimum, mean and maximum of instructed fixation period in each session among correctly performed trials. Each panel was an animal. Thick line was the ECDF of the mean fixation period in each session. Thin lines were the ECDFs of the minimum and maximum fixation periods, respectively.

**Figure S3.**
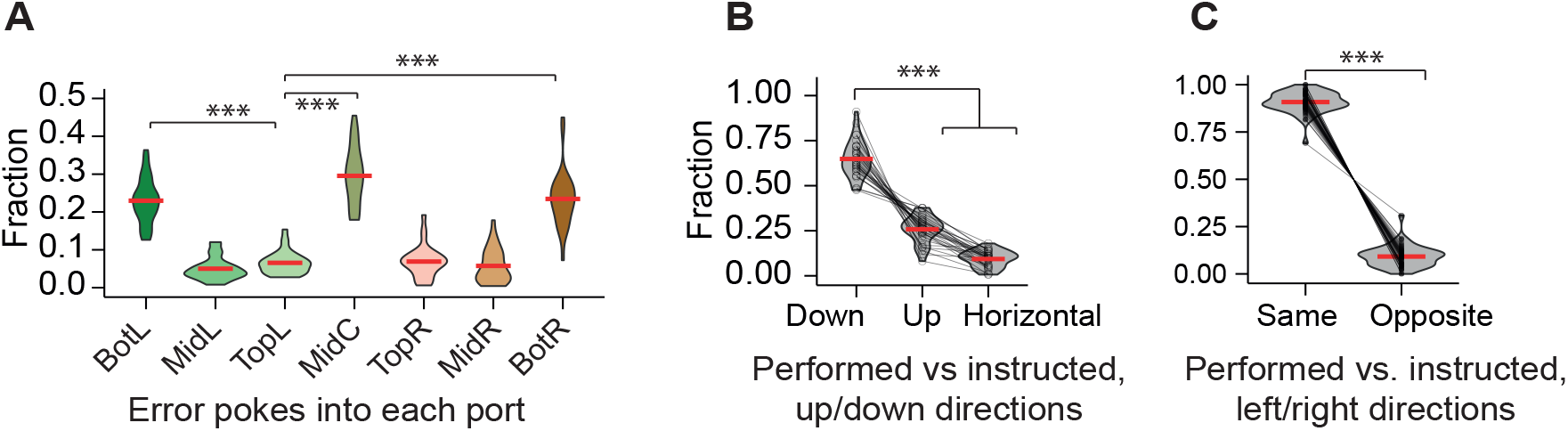
Error types of other rats in a task with the same temporal structure as the task in main result, but with different numbers of trial types. **A**. The proportion of error trials made into each port, grouped by subjects (n = 31). Rats prefer to make errors into the two lower ports and the central ports. **B**. The fraction of errors whose direction was lower, upper or horizontal compared to the instructed movement direction (31 subjects, 10934 trials). **C**. The fraction of errors whose direction was in the same side or opposite side to the instructed movement direction, among error trials that started from the central port (31 subjects, 10934 trials). ***, *p <* 0.001. Details specified in Table S2 for A and Table S3 for B and C.

**Figure S4.**
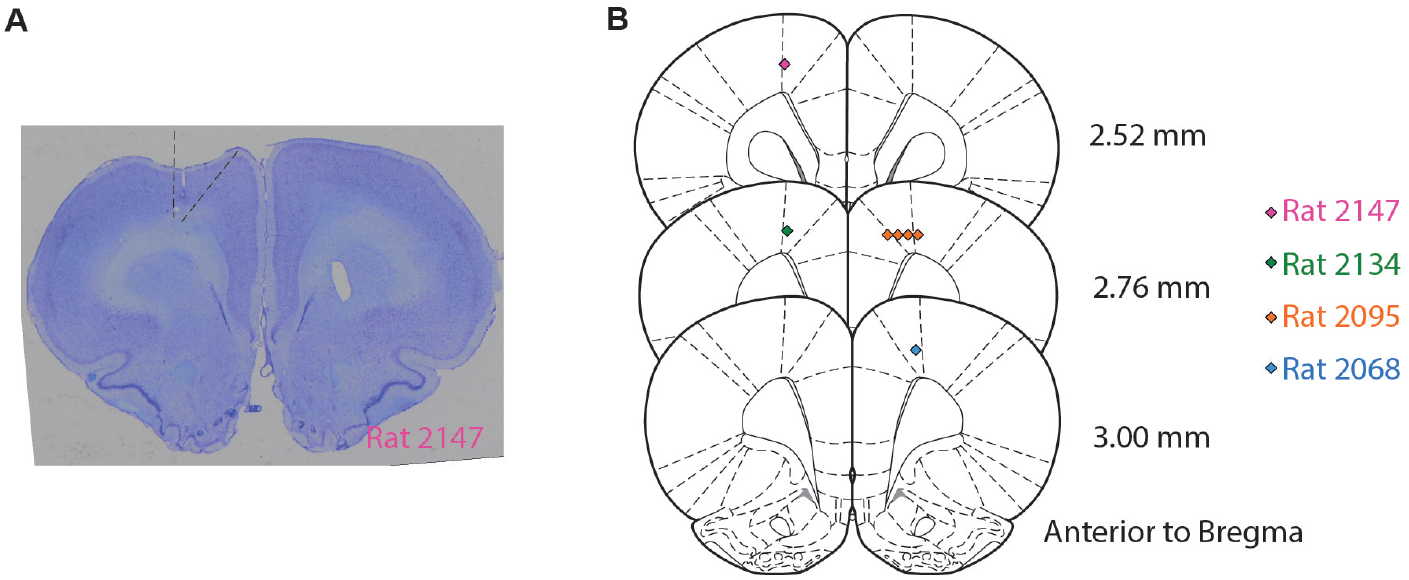
Histology. **A**. The coronal section of an example rat brain (2147) showing the placements of the silicon probes. Dashed lines indicate the estimated area of M2 in this brain section. **B**. The lesion marks matched to the coronal sections of the rat brain atlas(Paxinos and Watson, 2004). Lesions were made at the end of all the recording sessions with 200 uA for 3s relative to the GND. Colored marks indicate the lesion marks in **A**, and colors indicate subject ID. Recording sites in this manuscript varied across subjects in the anterior-posterior axis, ranging from AP 2.5 to 3.5 relative to the Bregma. These recording sites were within an head-orienting-related area based on previous anatomical, lesion, and microstimulation studies (Cowey and Ek, 1973, Crowne and Pathria, 1982, Leonard, 1969, Sinnamon and Galer, 1984), although more anterior compared to a previous electrophysiological study recording from the FOF literature where rats performed an auditory-guided 2AFC head orienting task (Erlich et al., 2011). Despite the variability in recording targets, the main findings in this manuscript were qualitatively consistent across subjects (Figure S6). Further experiments will be required to determine if there is a functional gradient over the anterior-posterior axis in the FOF.

**Figure S5.**
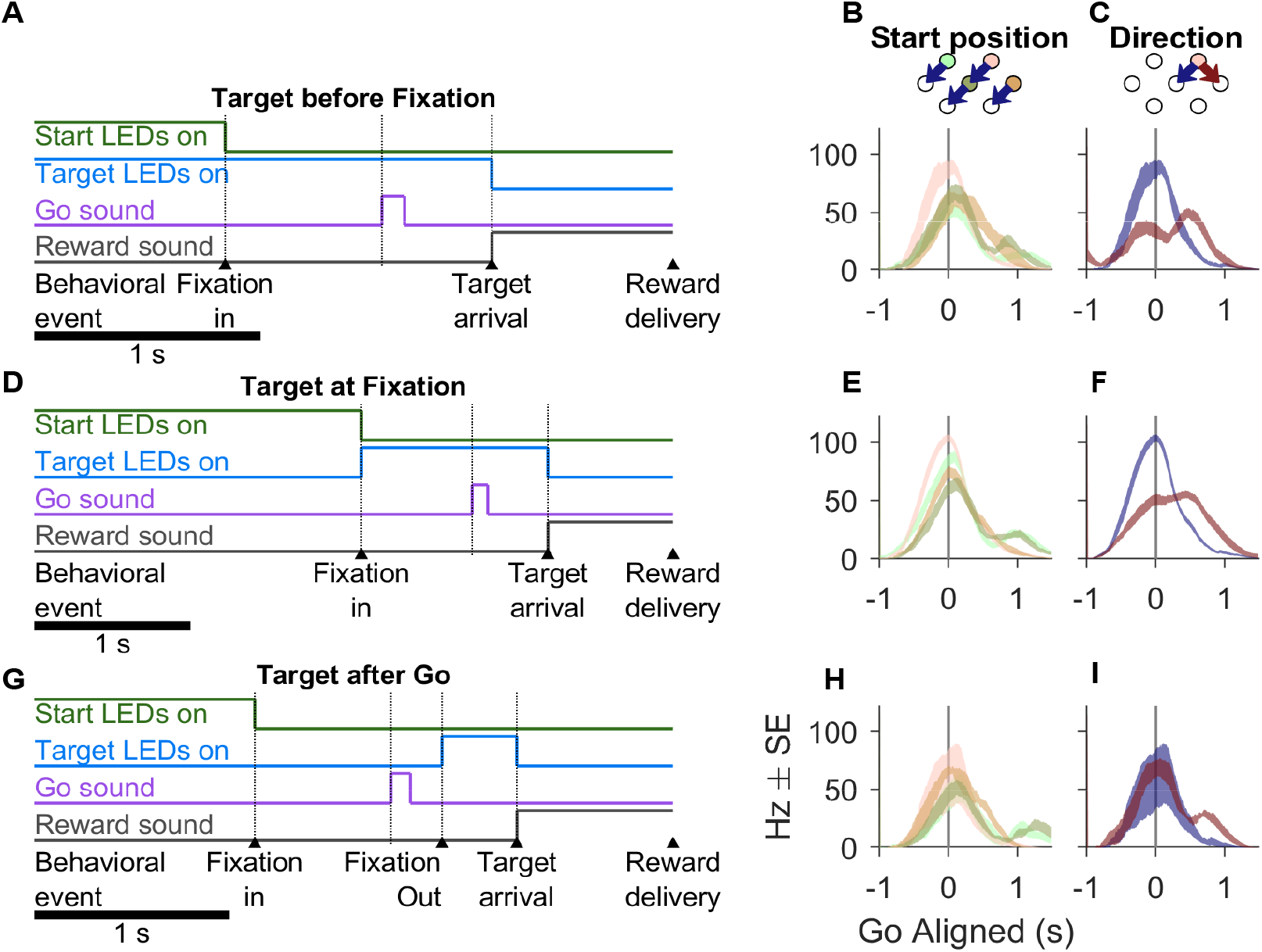
3 types of trial timelines in rat 2147, and the corresponding PETHs of an example neuron. In rat 2147, there were 2 types of sessions interleaved across days. In one session type, the trial timeline was similar to other animals, where the target LED was always illuminated when the rat fixated into the start port. In the other session type, there were 2 randomly interleaved types of timeline: “target before fixation” and “target after go”. The example neuron in B-C and H-I were recorded from the same session, and in E-F from another session. These units were considered as the same neuron based on the recording site, the waveform and the PETHs. **A**. The timeline of the **“target before fixation”** trial class. In these trials, the target LEDs and the start port LEDs illuminated at the same time, thus the target position information was available to the rat before fixation. **B**. The PETHs of an example neuron in “target before fixation” trials that were grouped by the start position and were into the same direction. **C**. The PETHs of the same example neuron as in B, but grouped by the movement direction and were from the same start position. **D-F** Similar to A-C, but for the **“target at fixation”** trials. In these trials, the target LEDs were illuminated when the animal poked into the start port. The neuron was selective to both the start position and the movement direction before the go sound. Note, that the neuron did not encode the movement direction earlier in C than in F, indicating that direction representation was related to the movement planning and not the visual cue. **G-I** Similar to A-C, but for the **“target after go”** trials. In these trials, the target LEDs illuminated after the animal has left the start port, which was detected by the IR sensors. Since the animal didn’t know the target port during fixation, the neuron was not selective to the movement direction before the go sound.

**Figure S6.**
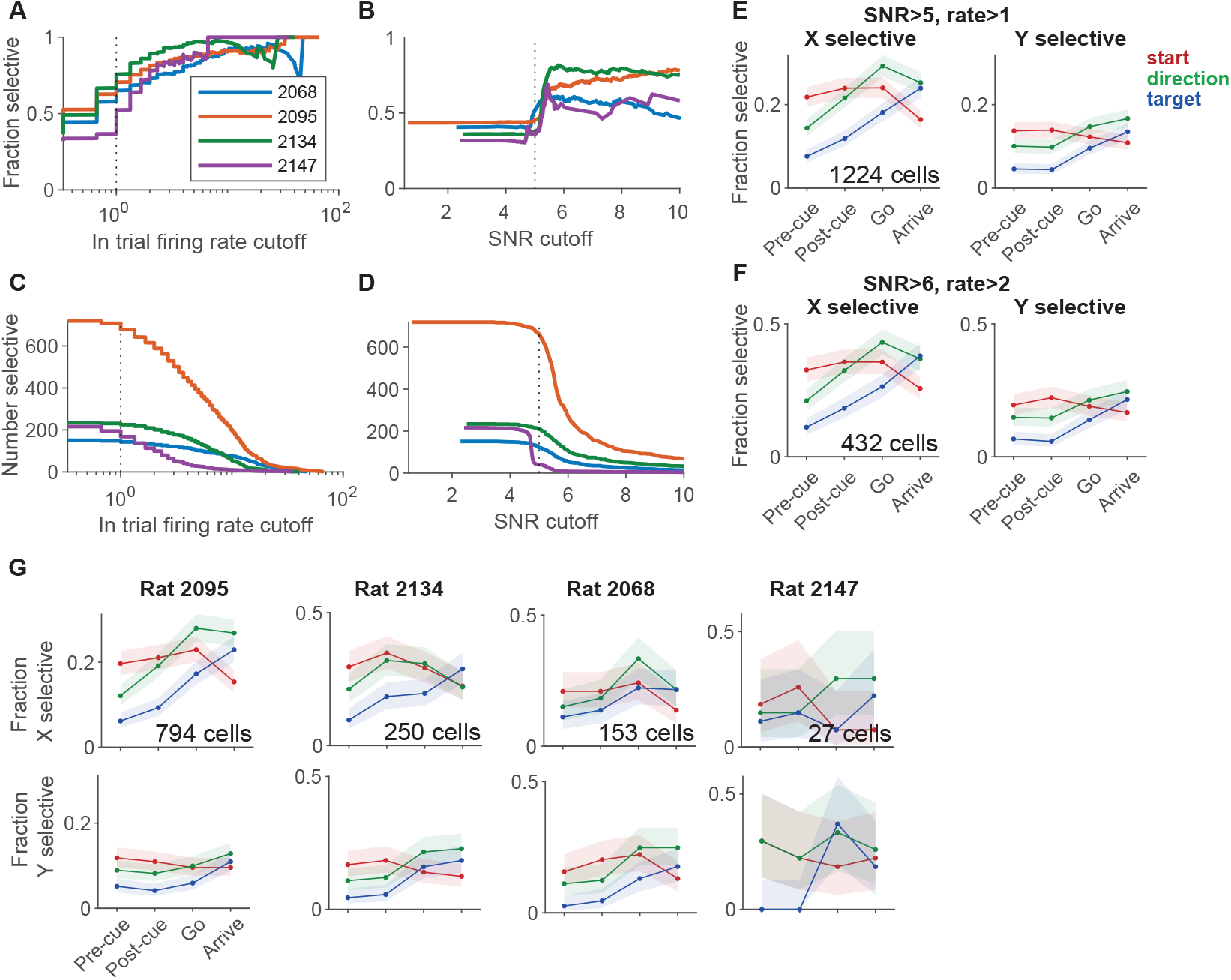
The sequential encoding of start position, direction and target position was consistent across subjects and single neuron selection criteria. In this figure, the selectivity of a neural response to a task variable was measured at each time point for the x and y coordinates separately using the area under the empirical ROC curve (the Wilcoxon-Mann-Whitney U-statistic). The p value was calculated by permutation test, where the null distribution was generated by shuffling the category labels 2000 times. **A**. The fraction of single neurons with spatial selectivity at different cutoff criteria for the in-trial firing rate. The signal-to-noise ratio cut-off was fixed at 5. A neuron is classified as spatially selective if the area under the empirical ROC curve is significantly larger than chance (*p <* 0.05, permutation test) for any one of the 3 spatial variables (start position, direction or target position) during any one of the 4 time windows (“pre-cue”, “post-cue”, “go” or “arrival”). **B**. The fraction of single neurons with spatial selectivity at different cutoff criteria for the signal-to-noise ratio (SNR) of the waveform. The cut-off criteria for in-trial firing rate was fixed at 1 Hz (*p <* 0.05, permutation test). **C**. The number of single neurons with spatial selectivity at different in-trial firing rate cutoff. **D**. The number of single neurons with spatial selectivity at different SNR cutoff. for the signal-to-noise ratio of the spike waveform. **E**. The fraction of neurons selective to the x coordinates (left panel) or y coordinates (right panel) of each spatial variable at the 4 time windows based on ROC analysis (*p <* 0.05, permutation test), when the SNR cutoff was 5 and in-trial firing rate cutoff was 1 Hz. Error bars were 95% confidence intervals of the binomial distribution. **F**. Similar to E, but the cut-off criteria is SNR *>* 6 and in-trial firing rate *>* 2. **G**. Similar to E, but for each subject.

**Figure S7.**
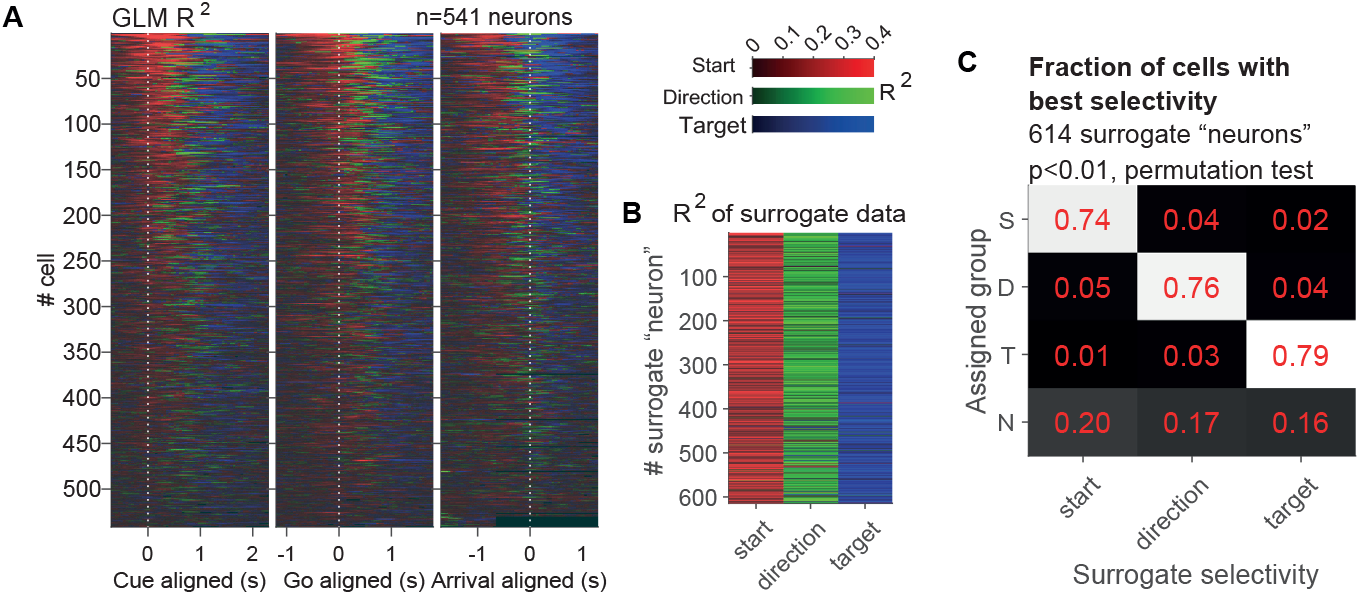
Validation of GLM analysis with surrogate data. **A**. The *R*^2^ of GLMs among neurons that were selective to at least one spatial variable in at least one of the 4 time windows (n = 541). *R*^2^s of GLMs are plotted over the right edge of the window (causal) of 300 ms with 50 ms steps. At each time window, only the GLM with the largest *R*^2^ is shown. Neurons are sorted by the total mass of the *R*^2^s of start position, direction and target position for the 3 alignments respectively. **B**. The *R*^2^s of GLMs in surrogate data with specific spatial selectivity. Each column was a group of surrogate data designed to have start, direction or target selectivity, and each row was a surrogate neuron. The color indicated the *R*^2^ of the model with the maximum *R*^2^, same as in Figure 3B. If the surrogate “neuron” had the largest *R*^2^ in the start position model, the corresponding cell was colored red, etc. Note, that the best model according to the *R*^2^s was visually consistent with the surrogate selectivity. **C**. Fraction of cells best selective to each spatial variable in surrogate data with specific spatial selectivity. There were 614 “surrogate neurons” in each column, designed to selective to the start position, the direction and the target position, respectively. The number in each row represented the fraction of “surrogate neurons” assigned as best selective to each model. S, start position model. D, direction model. T, target position model. N, non-selective to any one of the models. Surrogate data was generated to match the overall firing rates and the behavioral sessions of real neurons, but were known to selective to only one spatial variable (Methods). The best selectivity of a surrogate “neuron” was assigned in the same way as in Figure 3A. The assignment of “best selectivity” was correct in around 75% of surrogate “neurons”.

**Figure S8.**
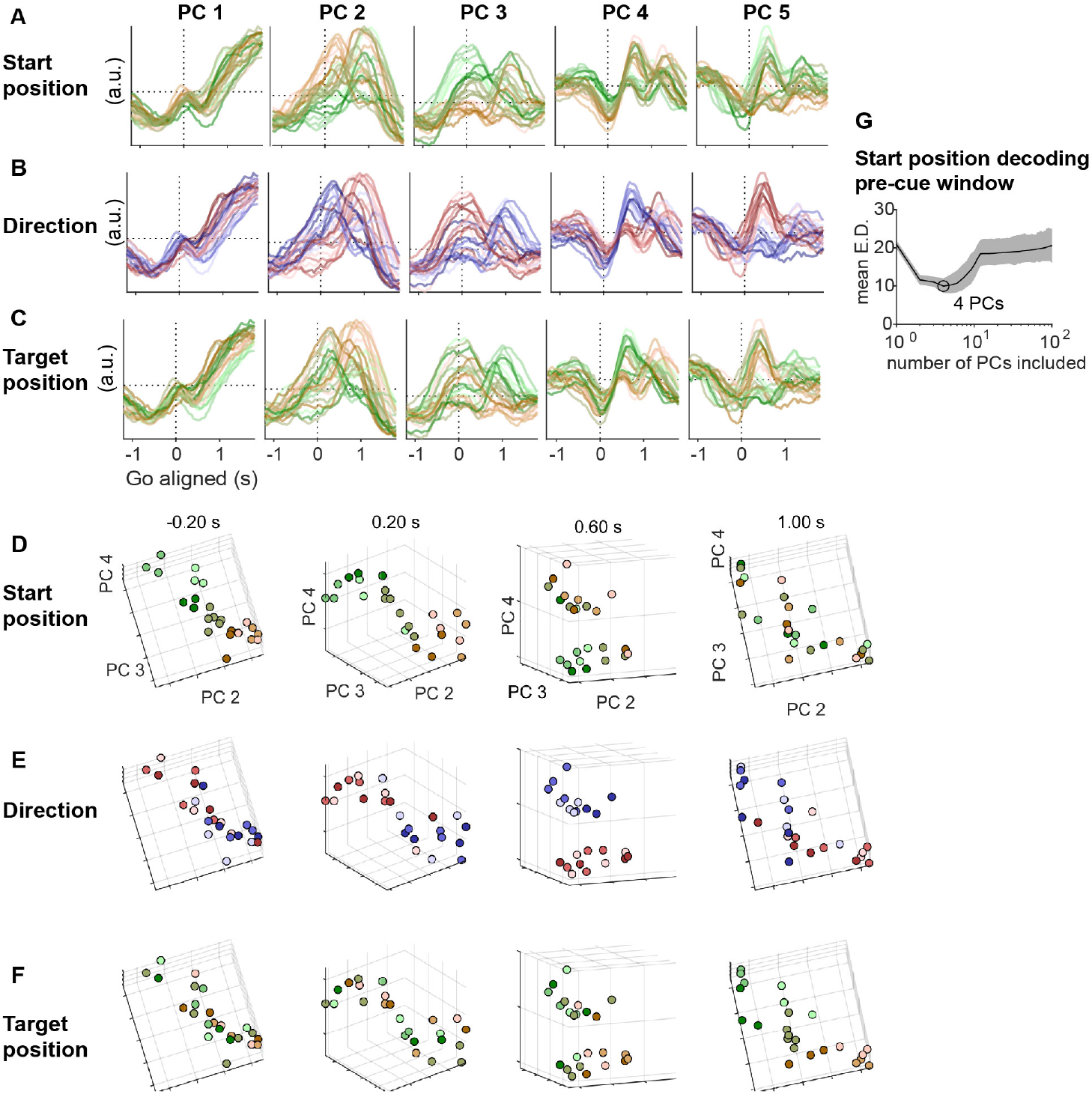
Principle components across time and trial types in the FOF population. **A**. The first 5 principle components of FOF population activity in each movement configuration across time aligned to the go sound, colored by the start position. The population was consisted of neurons with at least 8 trials in each of the 6 directions, 7 start positions and 7 target positions in “reference” sessions (771 neurons). Principle components were computed from a matrix of *N* by *C × T*, where *N* was the number of neurons, *C* was the number of trial configurations, and *T* was the number of time windows. **B-C**. Similar to A, but the PCs were colored by the direction and the target position, respectively. **D-F**. The coefficients of PC 2, PC 3 and PC 4 in each movement configuration at different time windows aligned to the go sound, colored by the start position (D), the direction (E) and the target position (F), respectively. The coefficients were rotated at each time window to aid visualization. In early stages of the trial (e.g. -0.20 s aligned to the go sound), trials with different start positions were well separated. As the movement initiates, direction became well separated in these PCs (0.20 s and 0.60 s). In later phases of the trial, the target position was well separated in these PCs (1.00 s). **G**.Number of principle components included versus pseudopopulation decoding accuracy. Mean error for start position decoding using spike count data in the pre-cue time window, among the same neurons and using same methods as in Figure 3 C, but including different numbers of principle components for the decoding. Thin line and shaded areas indicated the mean and the 5% and 95% intervals, as a result of re-sampling neurons with replacement for constructing the pseudopopulation.

**Figure S9.**
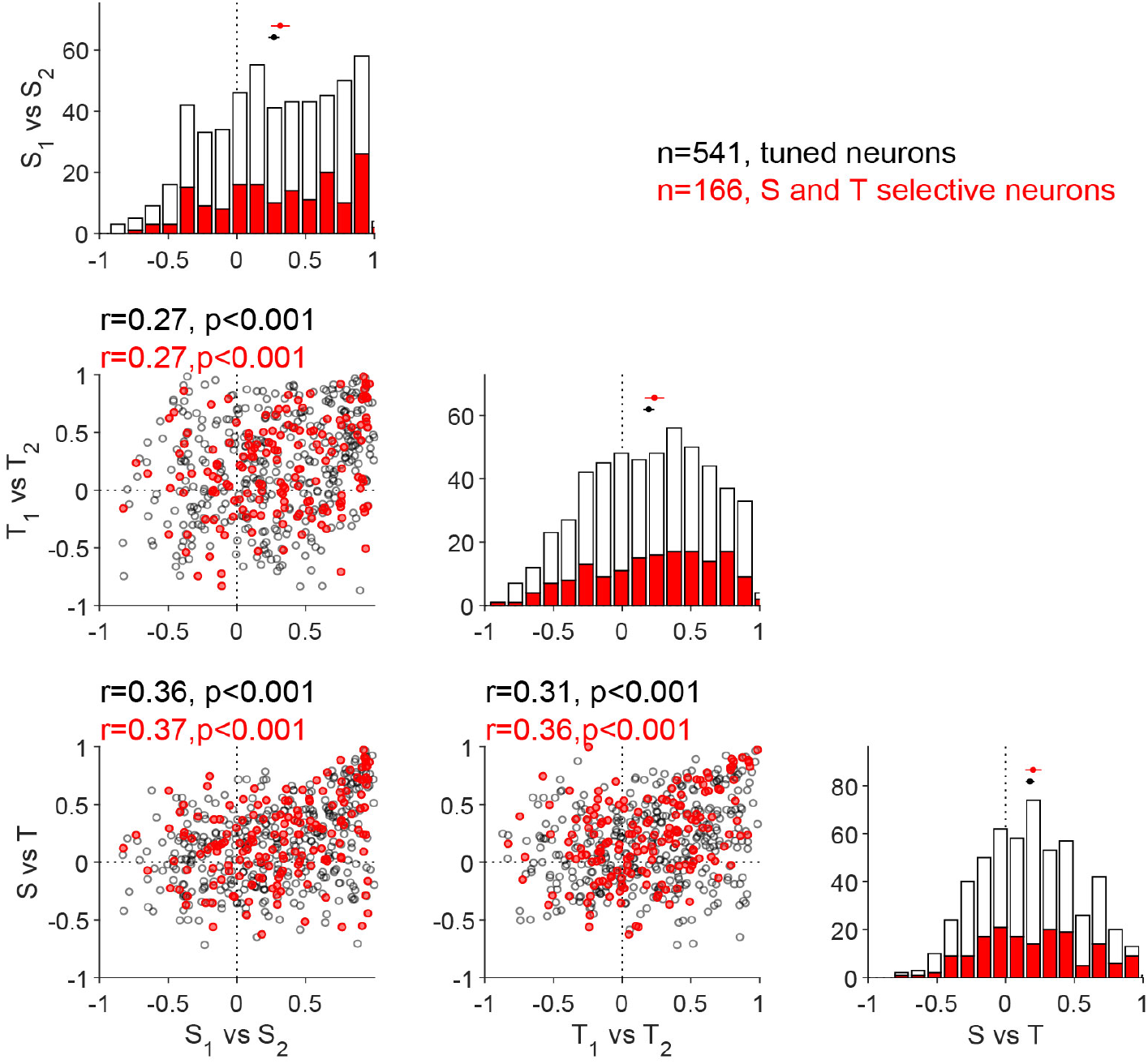
Start-target tuning correlation, start-start tuning correlation, and target-target tuning correlation were correlated. **Diagonal**. The three panels illustrate the number of neurons with different levels of start tuning correlation (at the pre-cue time window), target tuning correlation (at the arrival time window), or the start-target tuning correlation (pre-cue time window for start and arrival time window for target), respectively. The white bars indicate all the neurons with any spatial selectivity in any one of the “pre-cue”, “post-cue”, “go” or “arrival” time windows (*p <* 0.05, n= 808). The black bars indicate the neurons that were selective to the start position during the “pre-cue” window as well as the target position during the “arrival” window (*p <* 0.05, n=174). The black dot and horizontal line at the top of a panel indicate the mean *±*95% CI of the correlation coefficients among all the cells. **Scatter plots**. The tuning correlation of one pair of spatial variables versus the other. Black circles indicate all the spatially selective cells, matching white bars in the diagonal panels (n=541). Red circles indicate start and target selective cells, matching black bars in the diagonal panels (n=174). Texts at the top of each panel were the Spearman rank correlation coefficients of one *tuning correlation* versus the other, and text color matches the dot color. Start-start tuning correlation, target-target tuning correlation and start-target tuning correlation were significantly correlated with each other, i.e. neurons with consistent position selectivity tend to also be consistent between start and target selectivity.

**Figure S10.**
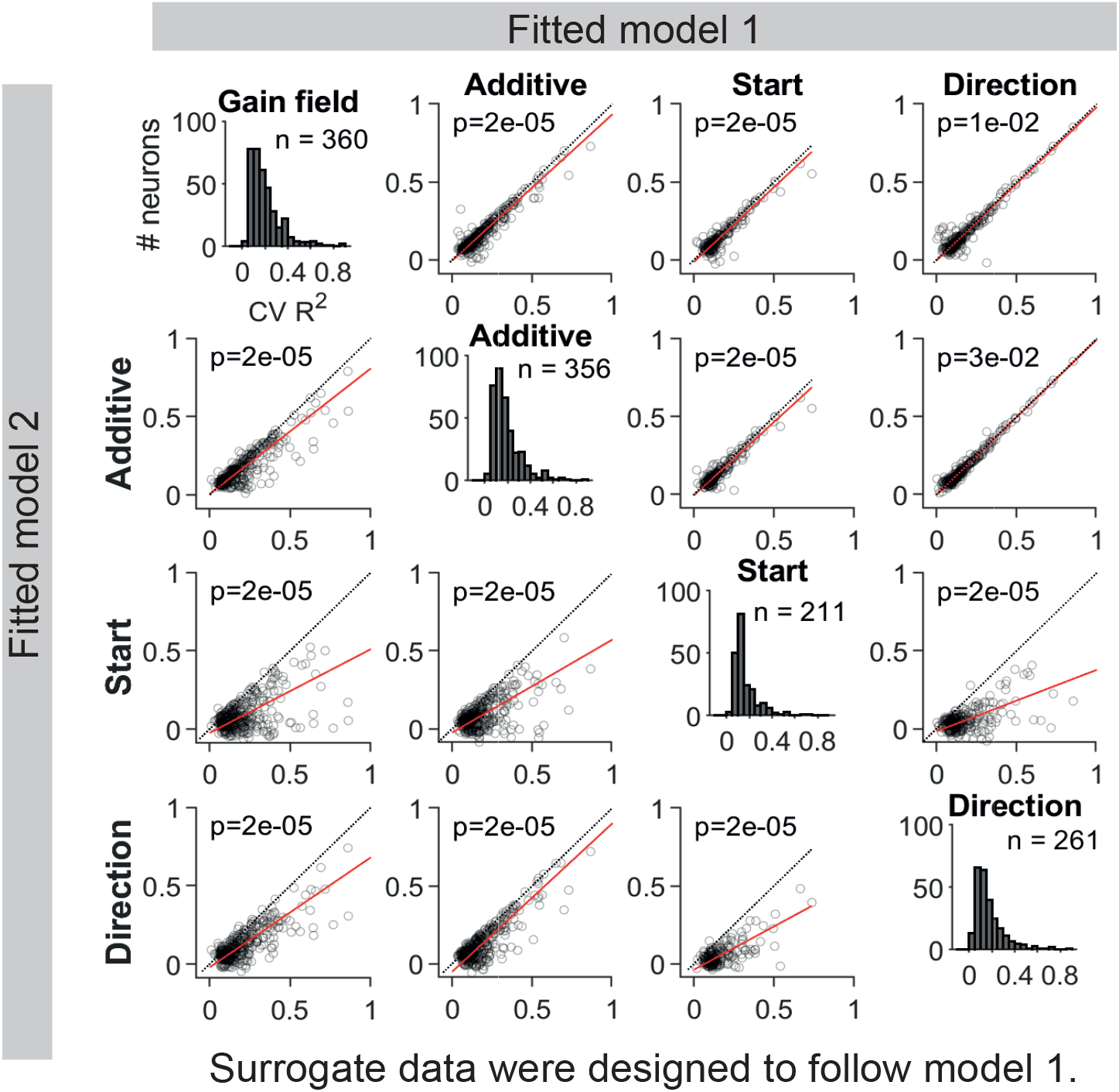
Detection of mixed selectivity with surrogate dataset. In all cases, our model comparison method correctly identified the generative model. **Off-diagonal scatter plots**. Each point is one surrogate “neuron” in each panel. The position on the x and y axes indicates the cross-validated *R*^2^s of one model versus the other. In each column of panels, the spike counts of surrogate “neurons” were designed to follow the same model as the fit model in the x axis. To generate the surrogate spike counts that followed the specific model, we fit the real neurons (n = 1224) to that model, and then generated Poisson distributed spike counts according to the predicted mean firing rates in each trial condition. We fit the surrogate spike counts to the spatial tuning models as in Figure 6, obtained 20-fold cross-validated *R*^2^s, and only plotted those surrogate “neurons” whose average of *R*^2^s in the 4 models were larger than 0.05. The number of included surrogate “neurons” in each surrogate dataset was indicated in the histograms. The *p* value in each panel was calculated with permutation test against the null hypothesis that the mean difference between the Fisher-z transformed *R*^2^s of the x axis model and the y axis model was not significantly different from zero (permuted 2000 times, n indicated in each panel). In each panel except the diagonal panels, the mean difference across surrogate “neurons” was significantly larger than 0, indicating that the model comparison with cross-validated *R*^2^s well captured the true functional form of spike count modulation. Notably, the method distinguished between the gain field and the additive model. **Histograms along the diagonal**. The distribution of cross-validated *R*^2^s for the 4 models, among surrogate “neurons” whose average of *R*^2^s in the 4 models were larger than 0.05 in each dataset.

**Figure S11.**
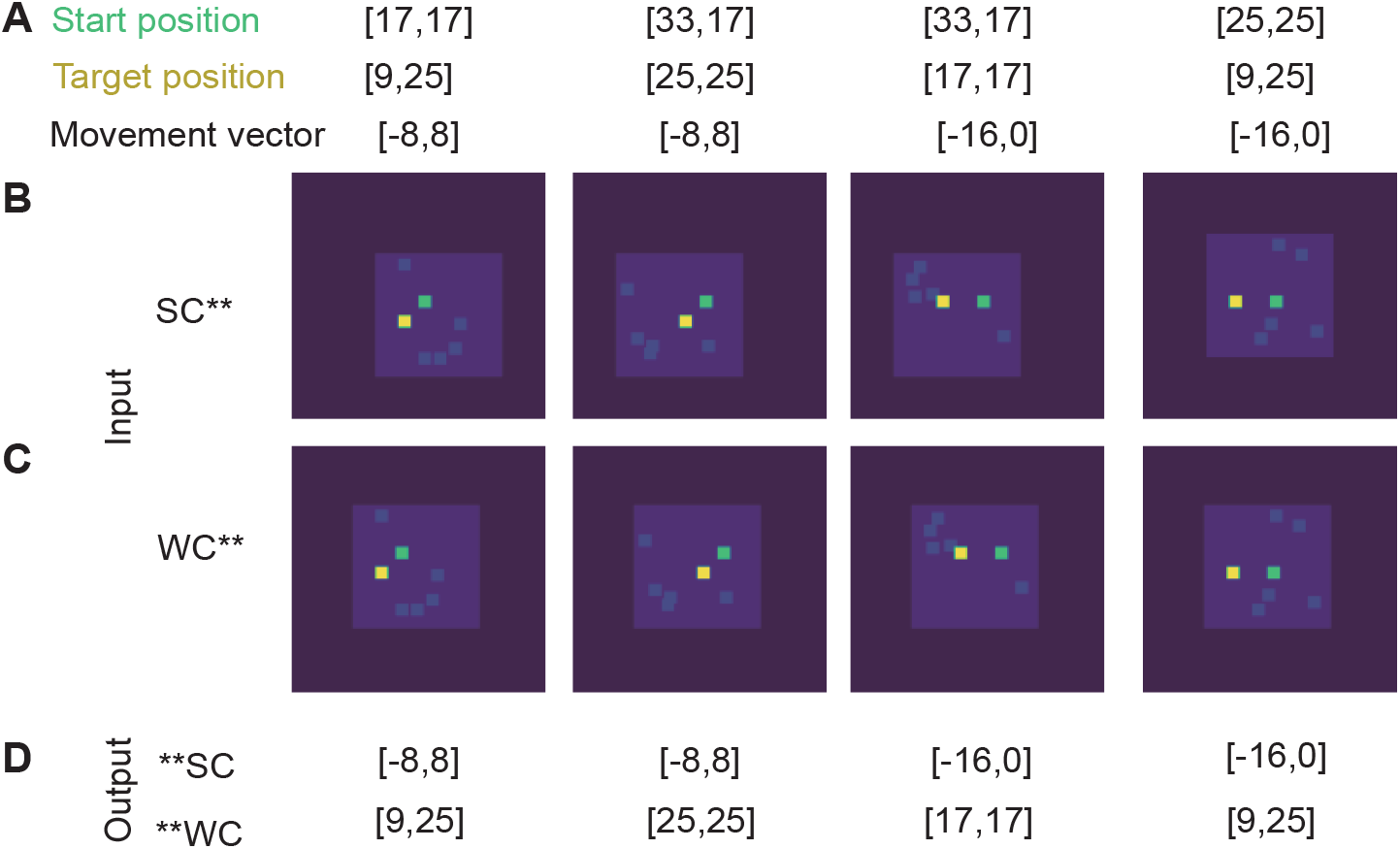
Examples of input and output configurations for recurrent network models. **A**. The coordinates of start position, target position and movement vector in a 100 *×* 100 pixel image for 4 example trials. The image itself denoted the visual frame, and the inner bounding box denoted the world frame. The start position coordinates and target position coordinates were relative to the bounding box. The movement vector was the difference between the target position and the start position. These coordinates were shown for the illustration purpose and were invisible to the models. **B**. An example input image for each trial when the input was in the self-centered coordinates (ego-*∗* networks). The green dot denoted the start position. The yellow dot denoted the target position. The light blue dots were distractors. The blue bounding box represented the world frame. Note, that the actual inputs were colorless and the elements were represented by intensities. **C**. An example input image for each trial when the input was in the world-centered coordinates (allo-*∗* networks). **D**. The output coordinates for the self-centered output (*∗*-ego) and world-centered output (*∗*-allo) models respectively. At the last time frame, the *∗*-ego models and the *∗*-allo models reported the movement vector or the target position coordinates respectively.

**Table S3.**
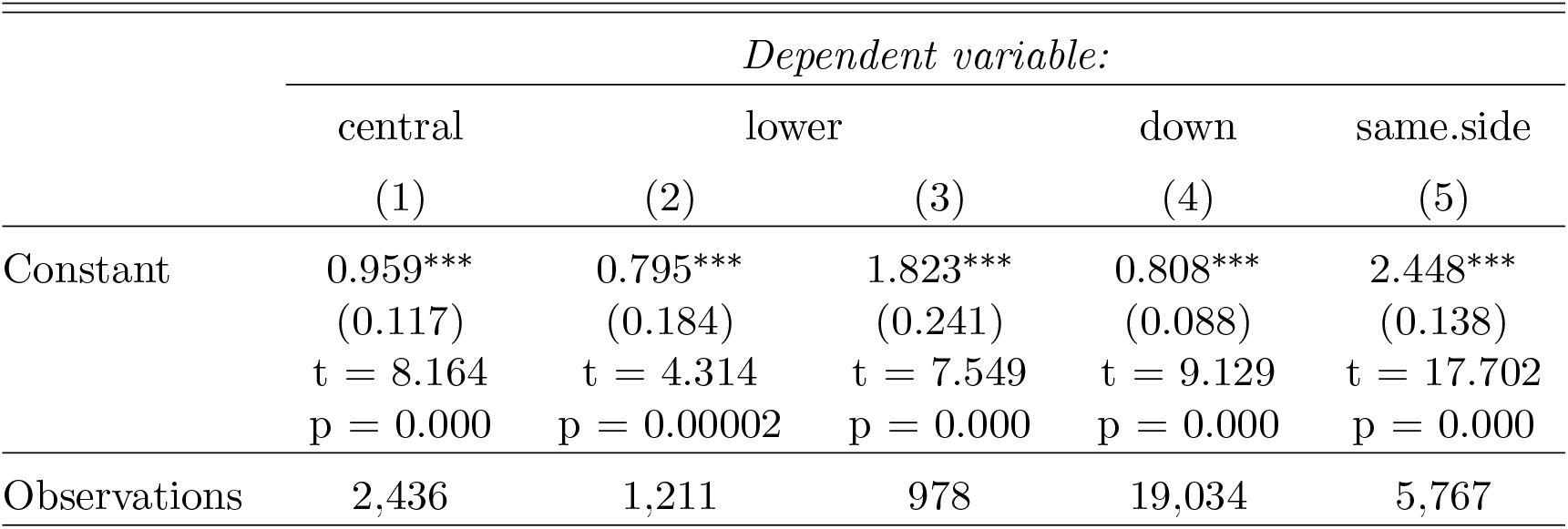
Bias in the fraction of error types quantified with logistic GLMMs. “Constant” was the fixed effect intercept term, *Estimate*(*SE*). Positive intercept term indicates that the frequency of the dependent variable being true was greater than 0.5. ^*∗*^p*<*0.1; ^*∗∗*^p*<*0.05; ^*∗∗∗*^p*<*0.01. These analyses only included “rand” group subjects. For (1)-(3), 21 subjects with long distance movements were include. For (4)-(5), 31 subjects included. (1)*central ∼*1 + (1|*subjid*), where *central* indicates that the trial landed in the central port. error trials with long-distance oblique movements were included. (2)*lower ∼*1 + (1|*subjid*), where *lower* indicates that the trial landed at the bottom port that was adjacent to the target port. Error trials with long-distance horizontal movements that landed at either the lower adjacent port or the central port were included. (3)*lower ∼*1 + (1|*subjid*),, where *lower* indicates that the trial landed at the bottom port that was adjacent to the target port. Error trials with long-distance horizontal movements that did not land at the central port were included. (4)*down ∼*1 + (1|*subjid*), where *down* indicates the error movement direction was vertically lower than the instructed movement direction. Error trials with short-distance movements were included. (5)*same*.*side ∼*1 + (1|*subjid*), where *same*.*side* indicates the error movement direction was horizontally the same as the instructed movement direction. Error trials short-distance movements starting from the center were included.

